# Harmala alkaloids regulate cell division planes in plants

**DOI:** 10.64898/2026.05.29.728883

**Authors:** Kamryn A. Diehl, Shayyaine Ocampo-Dallas, Olivia S. Hazelwood, Joh Demura-Devore, Fabiola Muro-Villanueva, Ryan S Nett, M. Arif Ashraf

**Affiliations:** Department of Botany, University of British Columbia, Vancouver, BC, V6T 1Z4 Canada; Department of Biology, Howard University, Washington, DC 20059, USA; Department of Molecular and Cellular Biology, Harvard University, Cambridge, MA, 02138, USA

## Abstract

Plants produce a vast diversity of specialized metabolites that function as chemical defenses against herbivores, pathogens, and competing plants. Many of these compounds also act as powerful tools for biological discovery, revealing fundamental cellular mechanisms through their effects on living systems. Among these metabolites, the harmala alkaloids from *Peganum harmala* (Syrian rue) possess cross-kingdom biological effects, including medicinal and neuroactive activity in humans, and allelopathic, growth-inhibiting effects on other plant species. However, the cellular processes in plants that are targeted by the harmala alkaloids are unknown. Here, we investigated the effects of the harmala alkaloids on plant growth and cell division using *Arabidopsis thaliana* as a model system. Of the harmala alkaloids, harmaline was identified as the most potent compound for root growth inhibition. Quantitative live cell imaging demonstrated that harmaline exposure causes progressive defects in cell division orientation and root cell morphology in a temporal manner. Furthermore, we identified harmaline-mediated phragmoplast orientation and morphology defects, pointing to a potential target related to phragmoplast guidance proteins. These findings position harmaline as a promising chemical probe for investigating the mechanisms that govern division plane positioning in plant cells and highlight a putative pathway by which harmala alkaloids exert allelopathic effects in competing plants.

**One sentence summary:** Harmala alkaloids regulate cell division plane as an allelopathic mechanism

## Introduction

Plant growth and development rely on the production of essential, primary metabolites like carbohydrates, proteins, lipids, and nucleic acids. Additionally, plants are well-known for their ability to synthesize a wide range of specialized metabolites for defense and chemical communication, which typically provide niche-specific ecological and evolutionary advantages. Specialized metabolites in plants can be broadly classified into alkaloids (caffeine, morphine, nicotine), terpenoids (paclitaxel, artemisinin, carotenoids), and phenolics (lignin, tannins, flavonoids), depending on their biosynthesis and chemical properties (Pichersky & Lewinsohn, 2011), and human civilizations have long explored the use of these molecules through thousands of years of cultural practice, curiosity-driven endeavor, and systematic medical research. Indeed, plants have served as a major source of specialized metabolites that are relevant to both medicine and agriculture (Craig & Newman, 2013). However, despite the importance of plants to human society, the biological roles of most specialized metabolites remain poorly understood, both in their native plant context and as tools for discovery (Bai et al., 2024). This represents a major knowledge gap in the study of plant specialized metabolism. Given the massive diversity of chemicals produced in the plant kingdom, it is likely that the exploration of native biological function for plant specialized metabolites will uncover novel mechanisms by which plants influence the biology of other organisms.

In this study, we sought to understand the reported allelopathic (plant growth-inhibiting) effects of *Peganum harmala* (Syrian rue), an understudied plant that exhibits notable adaptation to ecosystems with drought and high salt levels (Abbott et al., 2008). Among many other specialized metabolites, *P. harmala* is well-known for producing a group of β-carboline-containing molecules called the harmala alkaloids (harmol, harmalol, harmine, tetrahydroharmine or THH, and harmaline) (Figure 1A). These alkaloids accumulate to high levels in *P. harmala* roots and can affect the growth of other species, suggesting that they may contribute to the ability of *P. harmala* to outcompete nearby competitor plants (Herraiz et al., 2010). Interestingly, the five major harmala alkaloids differ in both concentration and spatial distribution throughout the plant, suggesting individual ecological roles for these alkaloids across tissue types and developmental stages (Hemmateenejed et al., 2006; Herraiz et al., 2010). For instance, harmaline is enriched in *P. harmala* seeds, whereas harmine is present at high levels in both seeds and roots (Herraiz et al., 2010; Shao et al., 2013). However, the genes required for biosynthesizing the harmala alkaloids are unclear, and thus little is known about the genetic basis for how spatial and temporal production of these alkaloids is controlled *in planta*.

**Figure 1:**
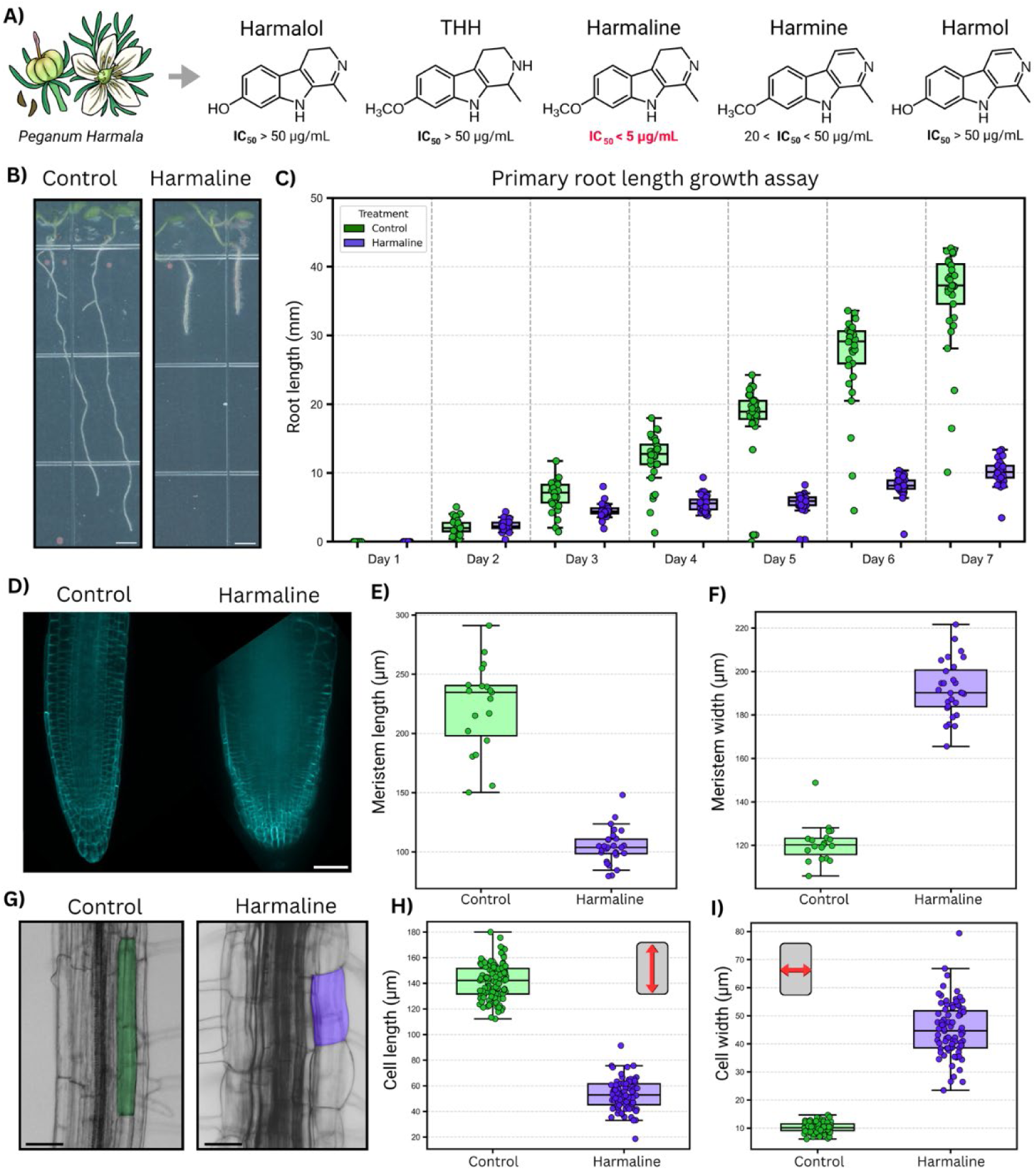
Harmaline inhibits root growth. **A)** List of main harmala alkaloids derived from *Peganum Harmala*. IC50 values were calculated from root growth assays in supplementary figure #. Harmaline has the lowest IC50 value and was thus used in this study. **B)** 5-days-old Arabidopsis seedling in absence and presence of 3μg/ml Harmaline. Scale bar = 3 mm. **C)** Quantification of root length from Figure 1A and Supplemental Figure 2. **D)** LTI6b-GFP root meristem in absence and presence of 3μg/ml Harmaline. Quantification of meristem length (E) and width (F) from Figure 1D. Scale bar = 50um.**G)** Epidermal cells of Col-0 root in absence and presence of 3μg/ml Harmaline. Scale bar = 50 um. Quantification of epidermal cell length (H) and width (I) from Figure 1G.

In support of a role for harmala alkaloids in allelopathy, extracts from *P. harmala* seeds, which contain high levels of harmaline, can inhibit root growth and germination in barley (Al-Mahmudy et al., 2024). Moreover, pure harmaline induces chlorosis and growth inhibition in *Arabidopsis thaliana* (Álvarez-Rodríguez et al., 2023) and alters cellular structures (Álvarez-Rodríguez et al., 2022). The allelopathic effect of harmala alkaloids also vary depending on the target species. For example, monocots such as wheat and ryegrass exhibit resistance to harmaline at concentrations that strongly inhibit growth in dicots like lettuce and amaranth (Shao et al., 2013). In contrast, harmine affects both monocots and dicots more broadly, without the same degree of specificity (Shao et al., 2013). Collectively, these studies highlight how harmala alkaloids could contribute to the allelopathic effects of *P. harmala* plants, but the cellular and molecular mechanisms of this activity have not been elucidated.

Here, we used the Arabidopsis root as a primary model for assessing how the harmala alkaloids impact root cell division and morphology. Our study determined that Arabidopsis roots are most sensitive to harmaline, with lower potency of root inhibition observed for the other tested harmala alkaloids. Furthermore, we found that harmaline affects Arabidopsis roots by altering the cell division plane, with a specific effect observed on phragmoplast orientation and cell elongation. Beyond exploring the impacts on cell division, we also found that physiologically relevant concentrations of harmaline affected the root growth of tobacco, camelina, and chia, while the root systems of crop plants like tomato, maize, and wheat were insensitive to harmaline. By using quantitative live cell imaging, we have highlighted a putative cellular mechanism for how harmaline exerts allelopathic activity, potentially through disruption of proper phragmoplast orientation during cell division. Given the variation of this phenotype among plant lineages, we expect that future comparative analyses among harmaline sensitive and insensitive plants will provide an important handle for deciphering the specific molecular targets of this alkaloid and its role in *P. harmala* allelopathy.

## Methods

### Plant materials

*A. thaliana* lines used for quantitative analyses included wild-type Columbia-0 (Col-0; ABRC stock CS6673), LTI6b-GFP (Kurup et al., 2005), Cytrap (Yin et al., 2014), proPCNA1::PCNA1-sGFP (Yokoyama et al., 2016), proTuB2::mScarlet-TuB2 (Schmidt-Marcec et al., 2023), pro35S::H2B-mRFP1 (Federici et al., 2012), TAN1-YFP (Rasmussen et al., 2011), YFP–RABA2a (Chow et al., 2008), and CDKA;1-mVenus/cdka;1 (Yang et al., 2020). Two double-marker lines, proPCNA1::PCNA1-sGFP × proTuB2::mScarlet-TuB2 and pro35S::H2B-mRFP1 × UBQ1pro::GFP-MBD, were generated in this study by genetic crossing. Segregating F2 and F3 seedlings were screened by confocal microscopy to identify seedlings expressing both fluorescent markers. Non-Arabidopsis species used in Figure 9 included tomato (Solanum lycopersicum; M82), wheat (*Triticum aestivum*; Ladoga), maize (*Zea mays*; B73), false flax (*Camelina sativa*; Varied), and black chia (*Salvia hispanica*; Compliments Naturally Simple).

### Plant growth conditions

*A. thaliana* lines expressing fluorescent markers were surface-sterilized and plated on half-strength Murashige and Skoog (½ MS) medium supplemented with 1% (w/v) sucrose. Seeds were sterilized by incubating in 70% ethanol for 10 minutes and washing three times with autoclaved MilliQ water. After plating, seeds were stratified in the dark at 4°C for 48h. Plates were then transferred to a growth chamber under continuous light at 22°C and grown vertically for three days. On day 3, seedlings were transferred either to control plates containing ½ MS with 1% sucrose or to treatment plates. For harmaline treatment, 3ug/mL harmaline plates were made by taking from a 5mg/mL harmaline stock and adding to autoclaved media that has cooled below ∼50°C but is still liquid. Seedlings were then returned to the growth chamber and maintained under continuous light at 22°C until imaging.

### Chemicals

Harmaline (Cayman chemical; Cat. no. #10995), Harmol (Cayman chemical; Cat. no. #33844), Tetrahydroharmine or THH (Cayman chemical; Cat. no. #14449), Harmalol (Cayman chemical; Cat. no. #35145), and Harmine (Cayman chemical; Cat. no. #10010324) were dissolved in organic solvent Dimethyl sulfoxide (DMSO) and mixed with ½ MS media for chemical treatment.

### Confocal microscopy

Cell outlines in control and harmaline-treated roots were visualized using either the plasma membrane marker LTI6b-GFP (Kurup et al., 2005) or the cell wall/cell outline dye propidium iodide. Cell-cycle progression was assessed using multiple fluorescent reporter lines. Cytrap was used to monitor S/G2 and G2/M transitions (Yin et al., 2014), while CDKA;1-mVenus/*cdka;1* was used to assess G1/S and G2/M transitions (Yang et al., 2020). A double fluorescent reporter line, proPCNA1::PCNA1-sGFP × proTuB2::mScarlet-TuB2, generated in this lab from proPCNA1::PCNA1-sGFP (Yokoyama et al., 2016) and proTuB2::mScarlet-TuB2 (Schmidt-Marcec et al., 2023), was used to distinguish G1/G2 from early S phase, and late S phase cell-cycle states. Live fluorescent imaging of *A. thaliana* root meristem and elongation zones shown in Figures 1 and 2 was performed using a Nikon Ti-E-PFS inverted spinning-disk confocal microscope equipped with a Yokogawa CSU-X1 spinning-disk unit, a four-line laser module, an Andor iXon 897 EMCCD camera, and a 20× objective in Howard University. Fluorescent images for all other figures were acquired at the UBC Bioimaging Facility (BIF) using an Evident IXplore SpinSR spinning-disk confocal system with a 40× objective lens. This system was equipped with a Yokogawa CSU-W1 spinning-disk unit, a 50 µm pinhole disk, and 405, 488, 514, 561, and 640 nm excitation lasers. Excitation at 488 nm, 514 nm, 561 nm, and 640 nm was used for GFP, YFP, RFP/mScarlet, and PI, respectively. Z-stacks were collected from the epidermal surface of the root inward to the deepest resolvable cell layers using a 0.3 µm step size and 25% laser intensity.

**Figure 2:**
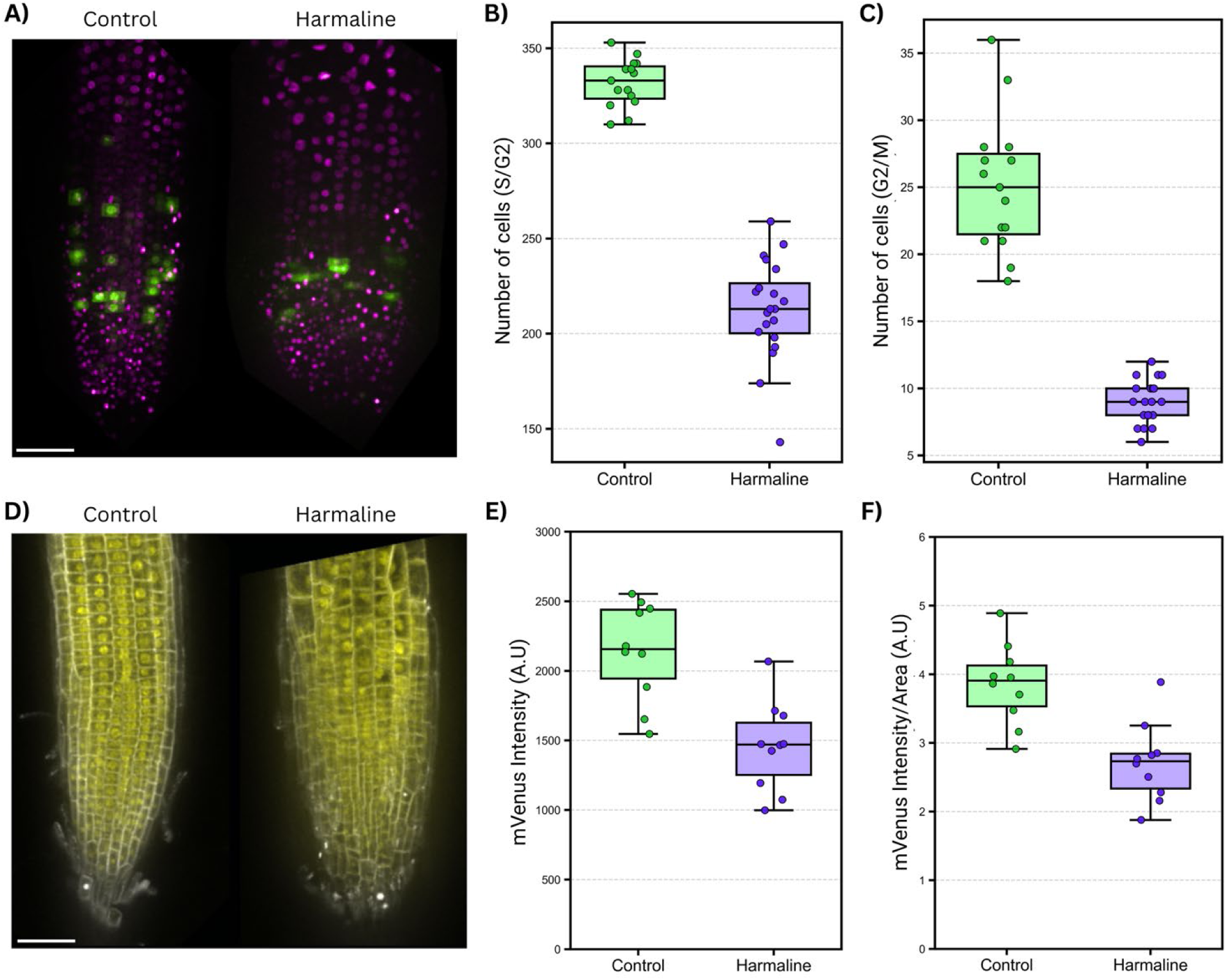
Cell cycle regulation in presence of harmaline. **A)** Cell cycle marker Cytrap (S/G2 and G2/M phases are indicated by magenta and green, respectively) in absence and presence of 3μg/ml Harmaline. Quantification of S/G2 **(B)** and G2/M **(C)** cell numbers from Figure 2A. **D)** Cyclin-dependent kinase, CDKA;1-mVenus (required for both G1 to S and G2 to M transitions) in absence and presence of 3μg/ml Harmaline. Quantification of mVenus intensity (A.U.) **(E)** and mVenus intensity (A.U.)/Area **(F)** from Figure 2D. Scale bar = 50 um.

### Scanning electron microscopy

*A. thaliana* Col-0 accession was grown on ½ Murashige and Skoog control and harmaline (2.5µg/mL) infused plates. At 7 days after germination, whole seedlings (n=20) were chemically fixed with osmium tetroxide, sputter coated with platinum and palladium using a Leica EM ACE600 Coater, and imaged with a Zeiss XB350 FIB-SEM.

### Quantification of confocal images

Z-stacks were examined sequentially through each root layer to quantify the number of normal and abnormal cells. Cells were classified as normal when the division plane was centrally positioned and showed no major morphological abnormalities. Cells were classified as abnormal when they exhibited one of the following division defects: triangular 1 or 2 divisions, in which small corner(s) cell displayed an incorrect division pattern; oblique divisions, in which the division plane was noticeably slanted rather than parallel to the expected orientation, except when such orientation appeared to avoid a junction with an adjacent cell wall; or vertical divisions, in which two adjacent cells showed normal horizontal division planes but the intervening cell displayed a vertical division plane. Only cells no larger than approximately twice the size of typical epidermal cells were included in the analysis, and root cap cells were excluded because they had already elongated. For root-swelling analysis, all epidermal cells located below the first visible root hair were measured using ImageJ/Fiji.

### Quantification of TAN1-YFP and RabA2a-GFP angles

TAN1-YFP and RabA2a-GFP localization angles were quantified from ImageJ measurements by extracting angular values from three defined positions within each cell: the apical (top) cell edge, the basal (bottom) cell edge, and the midpoint corresponding to the TAN1-YFP band or RabA2a-GFP plate. For each cell, these three measurements were treated as a single unit of analysis. To estimate the expected division plane orientation, a reference midpoint angle was calculated as the average of the top and bottom edge angles. Several derived metrics were then computed to assess TAN1-YFP band or RabA2a-GFP plate positioning relative to this reference. The one included in this paper was the aggregate “angle error” defined as the mean of the absolute deviations between the TAN1-YFP band or RabA2a-GFP plate and each edge. Absolute deviations were used where appropriate to capture magnitude independent of direction.

### Quantification of phragmoplast angle and curvature

Phragmoplast position and morphology were quantified from ImageJ/Fiji measurements using six measurements per cell collected in a fixed order using the line tool: (i) the top cell border angle, (ii) the bottom cell border angle, (iii) the phragmoplast angle, (iv) total cell length (perpendicular to the line used to measure i-iii), (v) the distance from one cell end to the phragmoplast midpoint, and (vi) a curvature angle measured with the ImageJ/Fiji angle tool if the phragmoplast was curved (midpoint line to the biggest angle the phragmoplast makes. This is visualized in Supplementary Figure 6.

To compare phragmoplast orientation with the local cell geometry, absolute values were taken for the top border, bottom border, and phragmoplast angles so that orientation could be assessed independent of sign. A local edge mean was then calculated as the average of the top and bottom border angles. Phragmoplast angular deviation was defined as the absolute difference between the phragmoplast angle and this local edge mean.

To quantify phragmoplast centrality within the cell, the measured end-to-midpoint distance was compared with total cell length. Distances were constrained to fall within the cell length, and a symmetric midpoint distance was calculated as the smaller of the two distances from the phragmoplast midpoint to either cell end. This value was also normalized to cell length to generate a midpoint fraction. Deviation from the true cell center was calculated as the absolute difference between the measured midpoint position and one-half of the total cell length.

### RNA-seq and analysis

RNA Isolation was performed using Arabidopsis Col-0 roots in control, harmine, and harmaline conditions using Qiagen RNeasy Plant Mini Kit (cat. no.: 74904). The extracted RNA concentration and quality was analyzed using Thermo Scientific NanoDrop One (cat. no.: 13-400-518). A total of 12 RNA samples (3 biological replicates of control along with 3 biological replicates harmine; and 3 biological replicates of control along with 3 biological replicates harmaline) were used. Poly-A enriched RNA-seq libraries were prepared by using Vazyme VAHTS Universal V10 RNA-seq Library Prep Kit and sequenced at 150-bp PE on an Element AVITI instrument by AmpSeq (Gaithersburg, MD, USA) (https://www.ampseq.com/).

RNA sequencing analyses were performed using the Fir high-performance computing cluster at Simon Fraser University. Raw paired-end sequencing reads were quality filtered and trimmed using Trim Galore with FastQC enabled (--paired --fastqc --quality 30 --gzip --length 50). Reads with Phred quality scores below 30 and reads shorter than 50 bp after trimming were removed. Filtered reads were aligned to the A. thaliana TAIR10.62 reference genome using HISAT2. Alignment output files were converted from SAM to BAM format using SAMtools. Transcript assembly and gene-level quantification were performed using StringTie with the TAIR10.62 genome annotation. Differential expression analysis was performed in R using the DESeq2 package based on gene read counts following a workflow adapted from Bahman’s analysis pipeline. Genes were considered differentially expressed if they exhibited a nominal *P* value < 0.05 and a fold change greater than 2-fold or less than 0.5-fold relative to controls. Variance stabilizing transformation (VST) implemented in DESeq2 was applied prior to principal component analysis (PCA), and sample clustering was visualized using the plotPCA function. Volcano plots were generated using the EnhancedVolcano package in R. Threshold lines corresponded to significance criteria of *P* < 0.05 and fold-change cutoffs of >2-fold or <0.5-fold. Gene Ontology (GO) enrichment analysis was performed using the clusterProfiler package with Arabidopsis annotations obtained through the org.At.tair.db package. Biological process GO terms were analyzed using the enrichGO function. Multiple-testing correction was performed using the Benjamini-Hochberg procedure, with an adjusted *P* value cutoff of 0.05 used to identify significantly enriched categories. Venn diagrams were generated from gene lists filtered using the same significance criteria (*P* < 0.05 and >2-fold or <0.5-fold change) to visualize overlap between differentially expressed gene sets.

### Statistical analysis

Raw data for each quantification were imported into JMP Pro 18 or analyzed using custom Python scripts in Jupyter Notebook. Python-based data analysis was performed using pandas for data import and organization, NumPy for numerical calculations, SciPy for statistical testing, and statsmodels for additional statistical analyses where appropriate. Data visualization was performed using Matplotlib, Seaborn, and Pathlib-based file handling for figure export. Statistical tests included Student’s *t*-tests, Mann-Whitney *U* tests, chi-square tests, and Fisher’s exact tests depending on the dataset structure and analysis goal. Data were plotted as individual measurements or replicate-level summaries, as indicated in each figure legend.

## Results

### Harmaline inhibits primary root growth

To investigate the root growth inhibitory effects of the major harmala alkaloids, we chose to work with plate-grown *Arabidopsis thaliana* as a tractable model system. Using this system, we first performed a dose response (5 µg/ml, 20 µg/ml, and 50 µg/ml) assay using harmine, harmaline, harmalol, harmol, and tetrahydroharmine (Figure 1A and Supplemental Figure 1). Among these five compounds tested, harmaline produced the strongest inhibitory effect on root elongation, with an IC_50_ value less than 5 µg/ml (Supplemental Figure 1). Harmine had the second highest potency, with an IC_50_ value of ∼20 µg/ml (Supplemental Figures 1 and 2). In contrast to harmaline and harmine, the other three compounds, harmalol, harmol, and tetrahydroharmine, only demonstrated clear root growth inhibition at the highest tested concentration (50 µg/ml). The dose response assay helped decipher the potency of major harmala alkaloids: harmaline > harmine > harmalol > harmol or tetrahydroharmine. As harmaline is the most potent compound among harmala alkaloids, we focused on subsequent analyses using 3 µg/ml harmaline.

We next performed a week-long time course assay using 3 µg/ml harmaline and measured the primary root growth of *A. thaliana* (Figure 1B and 1C; Supplemental Movie 1). The harmaline-induced root growth inhibition starts to appear on day 3 and this phenotype persists until day 7. The visible defects on root tissue were further confirmed by using scanning electron microscopy (SEM) imaging (Supplemental Figure 3). Because primary root growth is a combination of cell division and cell elongation (Rahman et al., 2007), we then examined the cellular cause of harmaline-induced root growth inhibition using live cell imaging. The plasma membrane localized marker, LTI6b-GFP, was used to measure the length and width of the meristem, which contains actively dividing cells (Figure 1D). We observed a dramatic decrease in meristem length and increase in meristem width (Figure 1E and 1F; Supplemental Movie 2). The reduction in meristem length suggests a decreased number of actively dividing cells. In contrast, the increased meristem width indicates the radial swelling of the root. To further understand the cell elongation and swelling, we observed the epidermal cells using light microscopy (Figure 1G). We found that under harmaline treatment, epidermal cell length was decreased, while cell width was increased (Figure 1H and 1I). Altogether, we found that harmaline seems to a) inhibit primary root growth by impacting both cell division and elongation, and b) induce radial swelling by increasing the individual cell width.

### Harmaline alters cell cycle regulation

The reduction in meristem size indirectly indicated that harmaline treatment might be causing a decrease in the number of actively dividing cells. To understand cell cycle activity, we performed live cell imaging using a dual-color cell cycle reporter, Cytrap, which indicates both S/G2 and G2/M phases simultaneously (Figure 2A) (Yin et al., 2014; Hazelwood & Ashraf, 2024). Quantifying cells at the S/G2 and G2/M phase indicated a reduction in the number of cells in both phases (Figure 2B and 2C). These results suggest an overall delay in cell cycle progression. We further confirmed the cell cycle progression using live cell imaging of CDKA1-mVenus/*cdka;1* counter-stained with propidium iodide for visualizing the cell borders. CDKA1 is a cyclin-dependent kinase which regulates both G1 to S and G2 to M transitions in *A. thaliana* (Cheng et al., 2015; Shimotohno et al., 2021; Iwakawa et al., 2006; Nowack et al., 2012; Hazelwood et al., 2026). With this method, we observed a reduction in CDKA1 signal after harmaline treatment, which further confirmed a slower progression of the cell cycle (Figure 2E and 2F).

### Harmaline affects the cell division plane in a temporal manner

We observed that control roots displayed the expected orderly pattern of transverse cell divisions, with newly formed cell plates positioned consistently parallel to existing cell walls (Figure 3A; Supplemental Movie 2). In contrast, harmaline-treated roots exhibited frequent aberrant divisions, including misplaced and irregularly oriented division planes that disrupted the normal cellular pattern of the root meristem (Figure 3A; Supplemental Movie 2). Quantification confirmed a strong increase in deviation in division plane angle in treated roots relative to controls, indicating that harmaline induces substantial defects in cell division plane orientation (Figure 3B). The quantitative live cell imaging for cell cycle markers (Cytrap and CDKA1-mVenus) and cell division plane (LTI6b-GFP) experiments were performed after 5 days of incubation with harmaline treatment, which led us to question the specificity of these phenotypes. To circumvent this issue, we performed quantitative live cell imaging, where 3-day-old seedlings were transferred to plates containing 3 µg/ml harmaline and imaged at 6h, 12h, 18h, 24h, and 48h time points post-transfer (Figure 3B). We started to observe the cell division plane defect as early as the 12h time point (3.97%), and the number of abnormal cell division events increased incrementally to 18h (4.57%), 24h (5.97%), and 48h (10.86%) (Figure 3C). We also observed a reduction of cell elongation and cell swelling starting from 6h, and this pattern became more obvious over the observation period (Supplemental Figure 4A-C). Additionally, we performed a correlation study to determine how cell elongation is related to the cell swelling phenotype. We only observed a positive correlation between cell elongation and cell swelling at the 12h and 24h time points (Supplemental Figure 4D). Among these observed phenotypes from the temporal data, the cell division plane defect correlates positively. To better understand the cell division plane defect, we classified the cell division plane defects into four categories: one triangle, two triangles, oblique, and vertical (Figure 3E). Based on these categories and our quantification, we found that the one and two triangle phenotypes increase over time and account for ∼50% of defects at 48h (Figure 3E).

**Figure 3.**
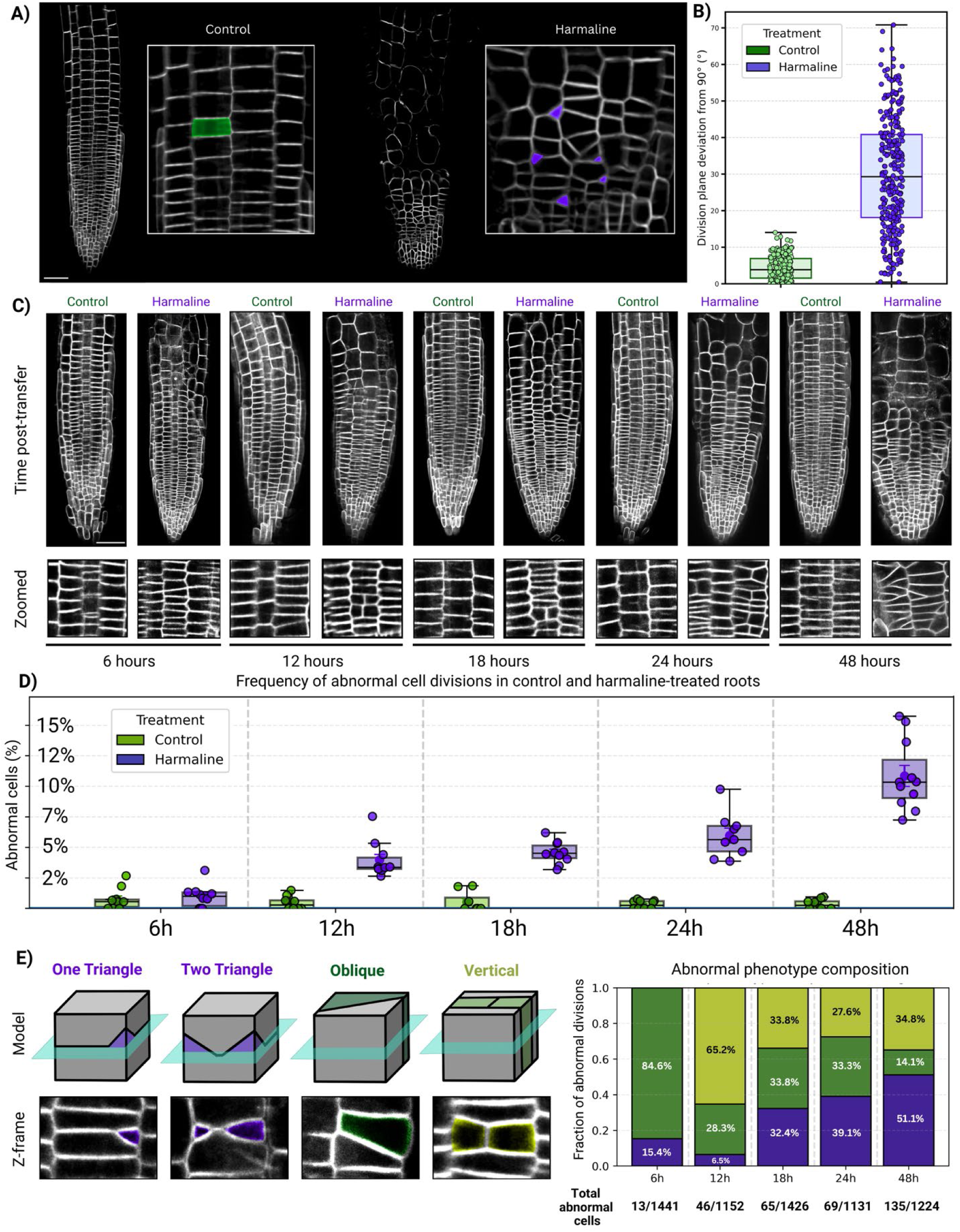
Harmaline induces time-dependent defects in root cell division. **A)** LTI6b-GFP root meristem in absence and presence of 3μg/ml Harmaline. Correctly formed division planes in control and aberrant cell division planes in presence of Harmaline are highlighted in cyan and magenta, respectively. Scale bar = 50 μm. **B)** Quantification of division plane positioning in LTI6b-GFP root meristems under control conditions and following treatment with 3 μg/mL harmaline. Division plane angles were transformed to represent deviation from the expected transverse orientation using angle−90|, where 0° indicates a perfectly positioned division plane and increasing values represent greater deviation from the normal orientation. Harmaline treatment increased the mean division plane deviation from 4.33 ± 3.32° (n = 190 cells) to 30.15 ± 15.64° (n = 280 cells) (mean ± SD; Welch’s t-test, p < 0.0001). **C)** Representative Spinning disk images of LTi6b-GFP Arabidopsis roots following transfer to 1% sucrose ½ MS media with or without 3ug/mL harmaline, imaged at 6, 12, 18, 24, and 48 h post-transfer. Harmaline-treated roots progressively display swollen, irregular cells and disorganized division planes compared to controls the same time after transfer. Bottom panels show zoomed views of the division zone for each time point. Scale bar = 50 μm. **D)** Frequency of abnormal cell divisions over time in control and harmaline-treated roots. Points represent individual biological replicates; box plots show median and interquartile range. Control roots maintained low abnormal division frequencies across all time points (6 h: 0.76 ± 0.27% SEM, n = 1489 cells; 12 h: 0.45 ± 0.17%, n = 1376 cells; 18 h: 0.53 ± 0.29%, n = 1113 cells; 24 h: 0.32 ± 0.11%, n = 1519 cells; 48 h: 0.35 ± 0.12%, n = 1598 cells). In contrast, harmaline treatment caused a progressive increase in abnormal divisions (6 h: 0.98 ± 0.30%, n = 1441 cells; 12 h: 3.97 ± 0.46%, n = 1152 cells; 18 h: 4.57 ± 0.29%, n = 1426 cells; 24 h: 5.97 ± 0.61%, n = 1131 cells; 48 h: 10.86 ± 0.86%, n = 1224 cells). Total replicates per condition was 10 each with cell counts ranging from 1,113-1,598 and 1,131-1,441 for harmaline treatments. **E)** Classification of abnormal division phenotypes, including single triangular cell plates, double triangular plates, oblique divisions, and vertical divisions. Illustrations show the green Z-plane intersecting the cell, and a representative image below from Lti6b-GFP. Stacked bars show the fraction of each phenotype over time among all abnormal cells (6 h: n = 13/1441; 12 h: 46/1152; 18 h: 65/1426; 24 h: 69/1131; 48 h: 135/1224), revealing a shift toward more severe division defects at later time points.

As the entire cell cycle requires 16-18h, we hypothesized that harmaline affects specific cell cycle stages instead of the entire cell cycle (Yin et al., 2014; Hazelwood & Ashraf, 2024). To track the individual cell cycle phases, we generated a cell cycle marker line containing *proPCNA1::PCNA1-sGFP* (indicates G1, late S, early S, and G2) with *proTuB2:mScarlet-TuB2* (preprophase band or PPB, spindle, and phragmoplast) (Hazelwood & Ashraf, 2024). We performed live cell imaging using this cell cycle line, and observed that harmaline-treated roots contained fewer cells overall than control roots, indicating a reduction in total cell production within the root meristem (Figure 4A). This is consistent with the reduced meristem size and lower abundance of cycling cells observed in Figure 1. However, we did not observe the difference in the proportional distribution of cells (interphase and mitosis) at different stages of the cell cycle (Figure 4B). To test whether harmaline preferentially altered progression through specific cell cycle phases, such as G1/G2, early S, and late S phases, we quantified the PCNA1-sGFP nuclear localization patterns (Figure 4C). Interestingly, the proportions of cells in G1/G2, early S, and late S did not differ significantly between control and harmaline-treated roots (Figure 4C). As we did not observe cell cycle phase-specific defects for pre-mitosis stages, we therefore next explored whether harmaline alters specific mitotic structures.

**Figure 4.**
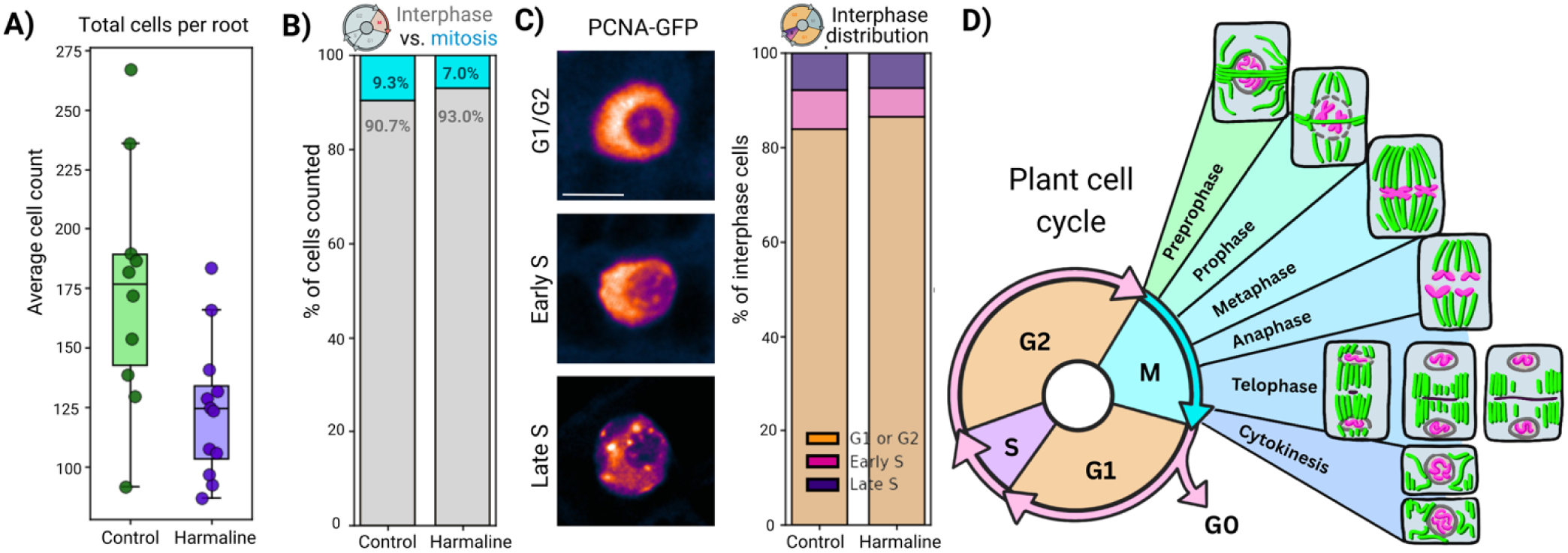
Harmaline treatment alters cell cycle distribution/reduces total cell counts without significantly shifting interphase composition. **A)** Total number of cells per root in control and harmaline-treated samples. Harmaline treatment resulted in a reduction in total cell counts (Control: n = 10 roots, 1749 cells total, mean = 174.9, SD = 50.84; Harmaline: n = 12 roots, 1492 cells total, mean = 124.33, SD = 29.20). This decrease was statistically significant (Welch’s t-test: t = 2.79, p = 0.0148; Mann-Whitney U = 98.0, p = 0.0134). B) Proportion of cells in interphase versus active mitosis. Harmaline treatment increased the proportion of interphase cells and decreased the proportion of actively dividing cells (Interphase: Control mean = 90.52%, SD = 1.88, n = 10 roots, 1749 cells; Harmaline mean = 93.09%, SD = 1.23, n = 12 roots, 1492 cells; Active mitosis: Control mean = 9.48%, SD = 1.88; Harmaline mean = 6.91%, SD = 1.23). These differences were statistically significant (Interphase: Welch’s t-test t = −3.72, p = 0.00206; Mann-Whitney U = 12.0, p = 0.00174; Active mitosis: Welch’s t-test t = 3.72, p = 0.00206; Mann-Whitney U = 108.0, p = 0.00174). **C)** Representative PCNA-GFP localization patterns used to classify interphase stages (G1/G2, early S, late S), and quantification of interphase distribution based on PCNA-GFP patterns. No significant differences were observed between treatments across stages (Control: n = 4 roots, 625 cells; Harmaline: n = 4 roots, 476 cells). Early S phase: Control mean = 8.15%, SD = 1.73; Harmaline mean = 6.15%, SD = 1.97 (Welch’s t-test: t = 1.53, p = 0.178). G1/G2: Control mean = 83.98%, SD = 4.65; Harmaline mean = 86.59%, SD = 2.59 (Welch’s t-test: t = −0.98, p = 0.376). Late S phase: Control mean = 7.87%, SD = 3.11; Harmaline mean = 7.27%, SD = 1.18 (Welch’s t-test: t = 0.36, p = 0.739). Scale bar = 5μm. **D)** Schematic overview of the plant cell cycle showing interphase (comprised of G1, S, G2) and mitotic stages (prophase through cytokinesis). Green represents microtubules and magenta represents DNA/chromosomes.

### Harmaline alters the phragmoplast orientation

To determine whether harmaline affects progression through specific stages of mitosis, we quantified the distribution of mitotic cells across preprophase band or PPB, metaphase, anaphase, early phragmoplast, and late phragmoplast stages (Figure 5; Supplemental Movies 3-5). Harmaline treatment altered this distribution in a stage-selective manner. The most striking difference was at the late phragmoplast stage, where harmaline-treated roots showed a clear accumulation of cells compared with controls. This enrichment suggests that cells either do not progress normally or take longer to undergo late cytokinesis under harmaline treatment. By contrast, metaphase, anaphase, and early phragmoplast populations did not differ significantly between control and harmaline-treated roots. This stage-specific enrichment of late phragmoplast cells aligns closely with the division plane phenotypes described in Figure 3.

**Figure 5.**
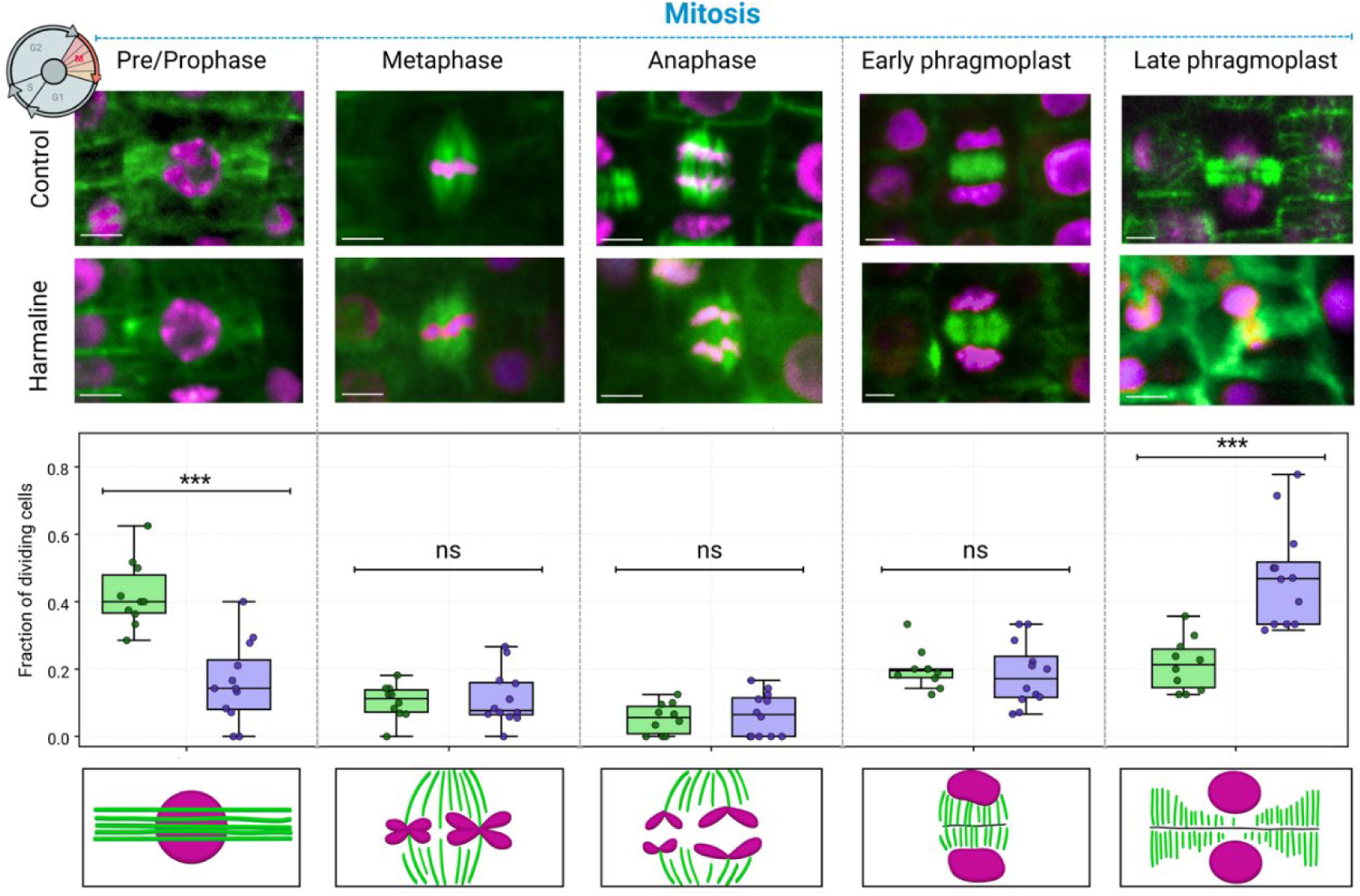
Harmaline-induced division defects arise in the root meristem and are associated with altered mitotic stage distribution. Representative images of dividing cells in control and 3 μg/mL harmaline-treated roots across mitotic stages (pre/prophase, metaphase, anaphase, early phragmoplast, and late phragmoplast). Microtubules are shown in green and DNA in magenta. Quantification of the fraction of dividing cells in each mitotic stage shows a significant redistribution under harmaline treatment. Harmaline-treated roots exhibit a strong reduction in pre/prophase cells (control mean = 0.422 ± 0.100 SD, n = 10 roots; harmaline mean = 0.160 ± 0.120 SD, n = 12 roots; Welch’s t = 5.58, p = 1.8 × 10⁻⁵; Mann-Whitney U = 114.0, p = 4.10 × 10⁻⁴) and a marked increase in late phragmoplast cells (control mean = 0.214 ± 0.079 SD; harmaline mean = 0.476 ± 0.151 SD; Welch’s t = −5.22, p = 6.8 × 10⁻⁵; Mann-Whitney U = 4.0, p = 2.47 × 10⁻⁴). In contrast, metaphase (p = 0.739), anaphase (p = 0.635), and early phragmoplast stages (p = 0.663) show no significant differences between treatments. An overall chi-square test of stage distribution confirms a significant shift in mitotic progression under harmaline treatment (χ² = 31.16, df = 4, p = 2.85 × 10⁻⁶), with aggregated counts showing depletion of early mitotic stages and accumulation at late cytokinesis (control vs harmaline mitotic stage counts: pre/prophase 70 vs 28; metaphase 18 vs 17; anaphase 9 vs 10; early phragmoplast 32 vs 28; late phragmoplast 34 vs 71). Scale bar = 5μm.

During mitotic stage classification, we noticed a prominent phragmoplast defect in harmaline-treated cells. We therefore classified late phragmoplasts as either normal or abnormal based on their morphology (Figure 6A). Control roots contained predominantly normal late phragmoplasts, which appeared relatively straight and symmetrically positioned across the division site (Figure 6A). Consistent with this, control late phragmoplasts were almost entirely normal (∼96%) (Figure 6A). In contrast, harmaline-treated roots showed a substantial increase in abnormal late phragmoplast morphologies (56%), including wavy, curved, and angled structures. This shift indicates that harmaline does not simply prolong late cytokinesis, but alters the geometry of the phragmoplast itself as it expands toward the cortex. In particular, curved and angled phragmoplasts were enriched under harmaline treatment, with the curved class population increasing from ∼4% in control roots to ∼29% under harmaline, and the angled class increasing from ∼9% to ∼14% (Figure 6A). These observations fit closely with the triangular and misoriented division plane phenotypes described in earlier experiments, and suggest that the abnormal cell plates arise from defective phragmoplast behavior during late cytokinesis rather than from a purely static positioning error. Additionally, we observed the phragmoplast deviation from the mid-plane of the cell in harmaline-treated roots compared to control (Figure 6B-D; Supplemental Figure 5). Together, these measurements demonstrate that harmaline disrupts both the positioning and geometry of the late phragmoplast, producing structures that are off-center and misaligned relative to the normal division plane.

**Figure 6:**
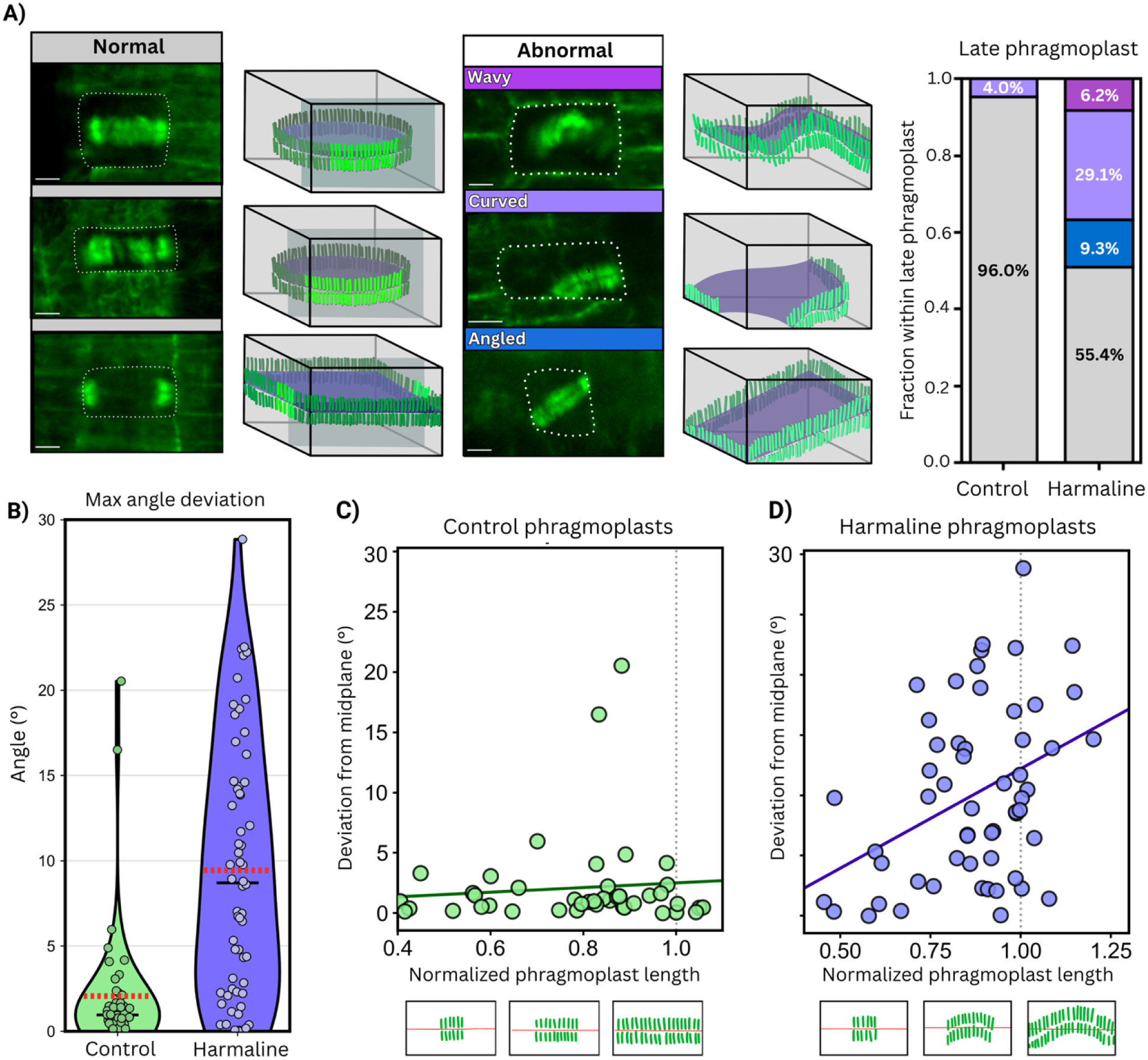
Harmaline treatment disrupts late phragmoplast morphology, leading to misoriented cell plates. **A)** Classification of late phragmoplast morphology into normal and abnormal categories (wavy, curved, angled), with representative images and 3D diagrams. Control cells are predominantly normal (mean = 0.96 ± 0.084), whereas harmaline-treated cells show a marked reduction in normal late phragmoplasts (mean = 0.55 ± 0.316; Welch’s t-test t = 3.92, p = 0.0027). Harmaline treatment increases the fraction of curved LPs (control mean = 0.04 ± 0.084; harmaline mean = 0.29 ± 0.233; t = -3.20, p = 0.0082), while increases in wavy (mean = 0.09 ± 0.168) and angled (mean = 0.06 ± 0.143) LPs are observed but not statistically significant across all replicates. **B-C)** Relationship between normalized phragmoplast length and maximum curvature angle. In control cells, curvature remains low and shows a weak negative correlation with length (Pearson r = −0.316, p = 0.032; n = 46 phragmoplasts). Harmaline-treated cells display elevated curvature across all lengths with no significant correlation (r = 0.093, p = 0.481; n = 60 phragmoplasts), suggesting a loss of length-dependent control of phragmoplast geometry. **D)** Combined data showing phragmoplasts in harmaline-treated cells exhibit greater curvature compared to control cells. Mean values are indicated by red dashed red lines. Harmaline treatment resulted in a substantial increase in curvature (Control: 2.07 ± 3.81, n = 46 phragmoplasts; Harmaline: 9.44 ± 7.40, n = 60 phragmoplasts). Statistical significance was assessed using Welch’s t-test (t = −6.65, p = 1.99 × 10⁻⁹). Scale bar = 5μm.

### Harmaline affects the new cell wall, not phragmoplast guidance, formation

It remained unclear whether these defects arise because harmaline was affecting the division plane specification or because they fail to correctly execute the division plane that has already been specified by proteins early in mitosis. We therefore compared TAN1-YFP, a marker protein required for phragmoplast guidance, with YFP-RABA2a, a marker of the newly forming cell plate, to distinguish between defects in phragmoplast guidance and new cell plate formation, respectively (Rasmussen et al., 2011; Chow et al., 2008). To determine whether harmaline alters phragmoplast guidance, we first examined TAN1-YFP localization at the cortical division site (Figure 7). In control cells, TAN1-YFP marked the expected division zone, and this localization pattern remained largely similar in harmaline-treated cells (Figure 7A). Even under harmaline treatment, TAN1 signal was still positioned at the expected cortical site, indicating that the cell retained the positional information needed to determine where cell division should occur. Consistent with this, the angle difference between TAN1 and the predicted division plane did not change significantly under harmaline treatment (Figure 7A). Together, these observations indicate that phragmoplast guidance remains largely intact and is unlikely to be the primary process disrupted by harmaline.

**Figure 7.**
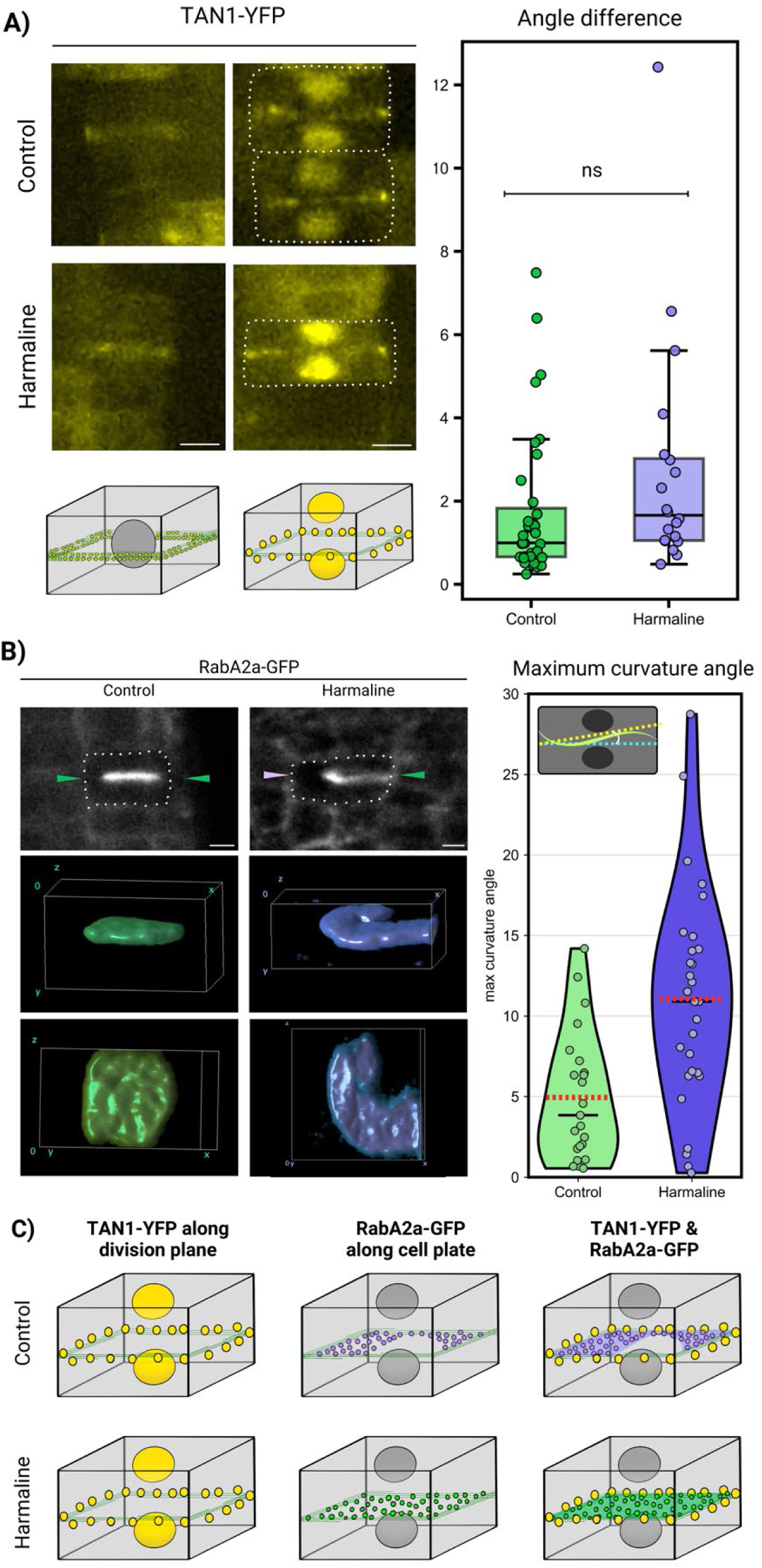
Harmaline alters phragmoplast and cell plate organization without strongly affecting division plane alignment. **A-B)** TAN1-YFP localization at the cortical division site in control and harmaline-treated roots. **A)** Representative images (top) of initial TAN1-YFP band and TAN1-YFP during cytokinesis, and schematic models (bottom). Quantification of angle deviation between TAN1-YFP signal and the expected division plane **B)** shows no significant difference between treatments (control: n = 35, mean = 1.67°, SD = 1.80; harmaline: n = 20, mean = 2.65°, SD = 2.84; Welch’s t = −1.38, p = 0.179), indicating that division plane specification is largely maintained under harmaline treatment. **C)** RABA2a-GFP localization at the forming cell plate in control and harmaline-treated roots. Representative single-plane images (top) and 3D reconstructions of those images (bottom) show a compact, planar cell plate structure in control cells, whereas harmaline-treated cells display irregular cell plate morphology. Quantification of maximum curvature angle gives a significant increase in curvature under harmaline treatment for both phragmoplasts (control: n = 46 phragmoplasts, mean = 2.07°, SD = 3.81; harmaline: n = 60 phragmoplasts, mean = 9.44°, SD = 7.40; Welch’s t = −6.65, p = 1.99 × 10⁻⁹) and cell plates (control: n = 23 cell plates, mean = 4.95°, SD = 3.97; harmaline: n = 29 cell plates, mean = 11.06°, SD = 6.78; Welch’s t = −4.06, p = 1.89 × 10⁻⁴). Quantification of maximum curvature angle gives a significant increase in curvature under harmaline treatment for cell plates (control: n = 23 cell plates, mean = 4.95°, SD = 3.97; harmaline: n = 29 cell plates, mean = 11.06°, SD = 6.78; Welch’s t = −4.06, p = 1.89 × 10⁻⁴). **E)** Summary schematics illustrating the relationship between TAN1-YFP (division plane marker) and RABA2a-GFP (cell plate marker). In control cells, the cell plate aligns closely with the division plane, whereas in harmaline-treated cells, the phragmoplast and cell plate are frequently curved or misdirected despite largely normal division plane specification. Scale bar = 5 μm.

In contrast, YFP-RABA2a revealed obvious abnormalities in the forming cell plate (Figure 7B; Supplemental Movies 6 and 7). In control cells, YFP-RABA2a-labeled cell plates appeared compact, smooth, and approximately planar (Figure 7B). Under harmaline treatment, however, cell plates appeared broader, less regular, and more poorly defined (Figure 7B). The 3D reconstructions made this difference especially clear, showing that harmaline-treated cell plates were not simply thicker, but also uneven and distorted in shape. These data indicate that the defect is not limited to where the division plane is positioned, but extends to how the cell plate is assembled and expanded within the cell (Figure 7B). Altogether, these experiments suggest that harmaline-induced aberrant phragmoplast orientation did not occur due to a defect in phragmoplast guidance (Figure 7C). Rather, the aberrant phragmoplast orientation leads to abnormal formation of new cells (Figure 7C). Moreover, the positional information for phragmoplast in the cell remains intact, but the newly forming phragmoplast fail to organize or assemble properly.

To gain initial insight on potential molecular targets of harmaline that may induce irregular cell division plane orientation, we performed a comparative transcriptomics analysis using harmaline (causes root growth inhibition and cell division plane defect) and harmine (causes root growth inhibition without cell division plane defect) with untreated roots as a control (Supplemental Figures 6-9, Supplemental tables 1-6). Gene expression analysis of selected cell cycle and division-associated genes did not reveal strong harmaline-specific transcriptional repression, and no major downregulated candidates emerged that could readily explain the observed division plane defects (Figure 8). This suggests that harmaline is unlikely to act primarily through broad transcriptional inhibition of known cytokinetic pathways. These findings are consistent with a model in which harmaline affects cellular function at a level downstream of transcription. Instead, harmaline may act through alternative mechanisms, including direct inhibition of protein function, disruption of protein localization or trafficking, altered protein stability, post-translational regulation, cytoskeletal dynamics, membrane-associated processes, or perturbation of signaling pathways required for proper division plane execution.

**Figure 8.**
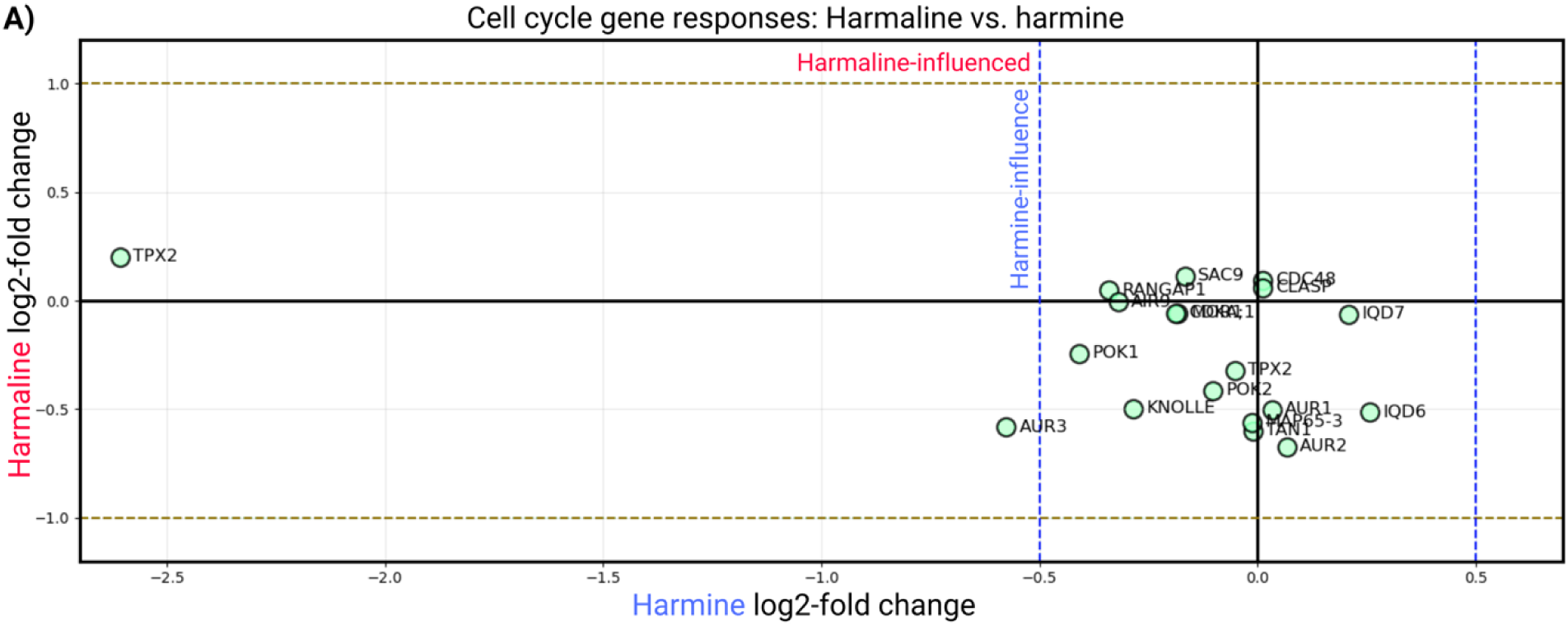
RNA-seq analysis of cell cycle and division-associated genes in response to harmaline and harmine treatment. **A)** Scatterplot comparing log2-fold expression changes of selected cell cycle and cell division-associated genes following harmaline (y-axis) and harmine (x-axis) treatment. Thresholds were set at ±1 for harmaline and ±0.5 for harmine to identify genes preferentially affected by harmaline treatment over harmine to determine whether proteins involved in division plane establishment/cytokinesis showed selective transcriptional responses that could explain why harmaline induces division plane defects whereas harmine does not. Genes examined included cyclins and cell cycle regulators (CYCB1;1, CYCB2;1, CDKA;1, CDKB1;1, CDKB2;1, CDC2, CDC48), cytokinesis and cell plate proteins (KNOLLE/SYP111, MAP65-3, PLE), division plane and microtubule-associated proteins (TAN1/ATN, POK1, POK2, TPX2, AUR1, AUR2, AUR3, RANGAP1, IQD6, IQD7, CLASP, MOR1, AIR9), and additional regulatory proteins (SAC9). No genes met the predefined criteria for strong harmaline-specific responses. Two genes, TPX2 and AUR3, showed stronger expression changes in response to harmine than harmaline.

### Harmaline has distinct allelopathic effects across the plant kingdom

Our quantitative live cell imaging data suggests that harmaline affects the phragmoplast orientation and consequently, the late cytokinetic steps in dividing *A. thaliana* root cells. Because cell division machineries are highly conserved across the plant kingdom (Jürgens, 2005), we next asked how broadly this response is conserved across plant species. Previously, it had been shown that harmine and harmaline had inhibitory effects on dicots (lettuce and amaranth), with monocot species (wheat and ryegrass) being more resistant (Shao et al., 2013). To expand on these previous results and examine the broader distribution of harmaline sensitivity, we compared primary root growth across a phylogenetically diverse panel of plant species, including *Zea mays* (maize), *Triticum aestivum* (wheat), *Camelina sativa* (camelina), *Salvia hispanica* (black chia), *Solanum lycopersicum* (tomato), and *Nicotiana tabacum* (tobacco) (Figure 9, Supplemental Movies 8-10). Among the species tested, *A. thaliana*, *N. tabacum*, *S. hispanica*, and *C. sativa* showed particularly strong sensitivity to harmaline, with clear decreases in root length accompanied by visible morphological changes relative to control seedlings (Figure 9, Supplemental Movies 8-10). In contrast, *Z. mays*, *T. aestivum*, and *S. lycopersicum* showed little to no visible difference between control and harmaline-treated seedlings (Figure 9, Supplemental Movies 8-10). Furthermore, we tested the effects of harmaline on *N. tabacum* and *C. sativa* root growth at the cellular level (Figure 10), which revealed an altered division plane and cell swelling phenotype in harmaline-treated roots (Figure 10). In a mechanistic context, the conserved cellular phenotype (cell division plane and cell swelling) of harmaline in three distinct plant organisms’ root systems highlights a likely conserved allelopathic mechanism of this molecule at the cellular level. More broadly, because our study highlighted that many crop plants (e.g. maize, wheat, and tomato) are resistant to harmaline, deciphering the molecular target and resistance-related genes for this alkaloid may provide an excellent opportunity for identifying novel herbicidal mechanisms of action.

**Figure 9.**
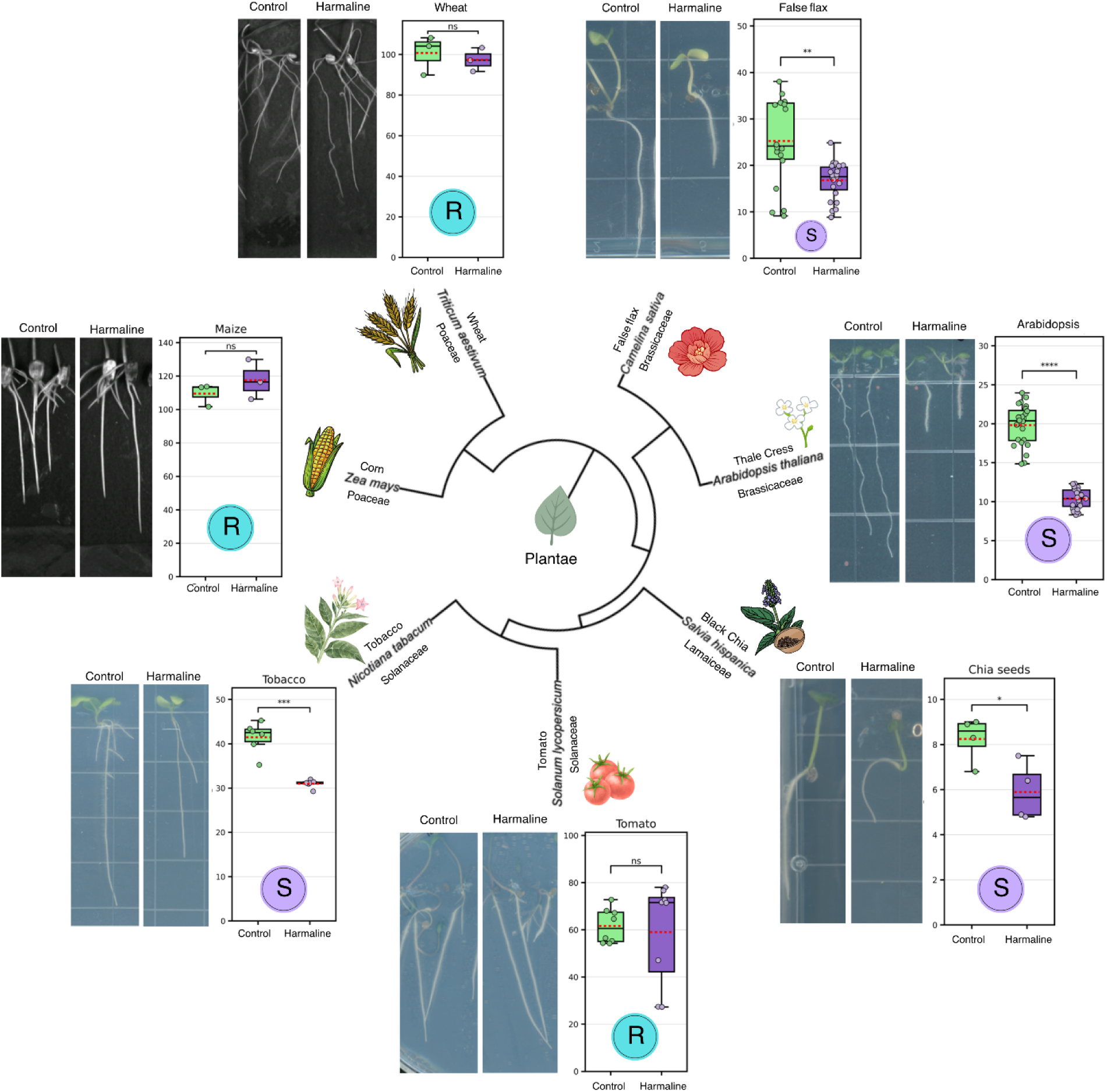
Harmaline elicits conserved growth inhibition with species-specific sensitivity across diverse plant lineages. Representative images and quantitative analyses of root growth in multiple plant species grown under control and harmaline treatment are shown alongside a phylogenetic framework. Harmaline significantly reduced root growth in Arabidopsis thaliana (Control: 19.83 ± 2.59; Harmaline: 10.38 ± 1.26; Welch’s t = 16.06, p < 0.0001, ****), Nicotiana tabacum (Control: 41.48 ± 3.52; Harmaline: 30.99 ± 0.91; t = 7.07, p = 0.0005, ***), Camelina sativa (Control: 25.26 ± 9.42; Harmaline: 16.81 ± 3.95; t = 3.57, p = 0.0018, **), and Salvia hispanica (chia) (Control: 8.25 ± 1.01; Harmaline: 5.90 ± 1.29; t = 2.86, p = 0.0307, *). In contrast, no significant differences were observed in Solanum lycopersicum (tomato) (Control: 61.65 ± 7.35; Harmaline: 59.05 ± 21.76; t = 0.32, p = 0.7571, ns), Zea mays (maize) (Control: 109.56 ± 6.82; Harmaline: 117.51 ± 11.94; t = −1.00, p = 0.3867, ns), or Triticum aestivum (wheat) (Control: 100.71 ± 9.61; Harmaline: 97.36 ± 5.83; t = 0.52, p = 0.6383, ns).

**Figure 10.**
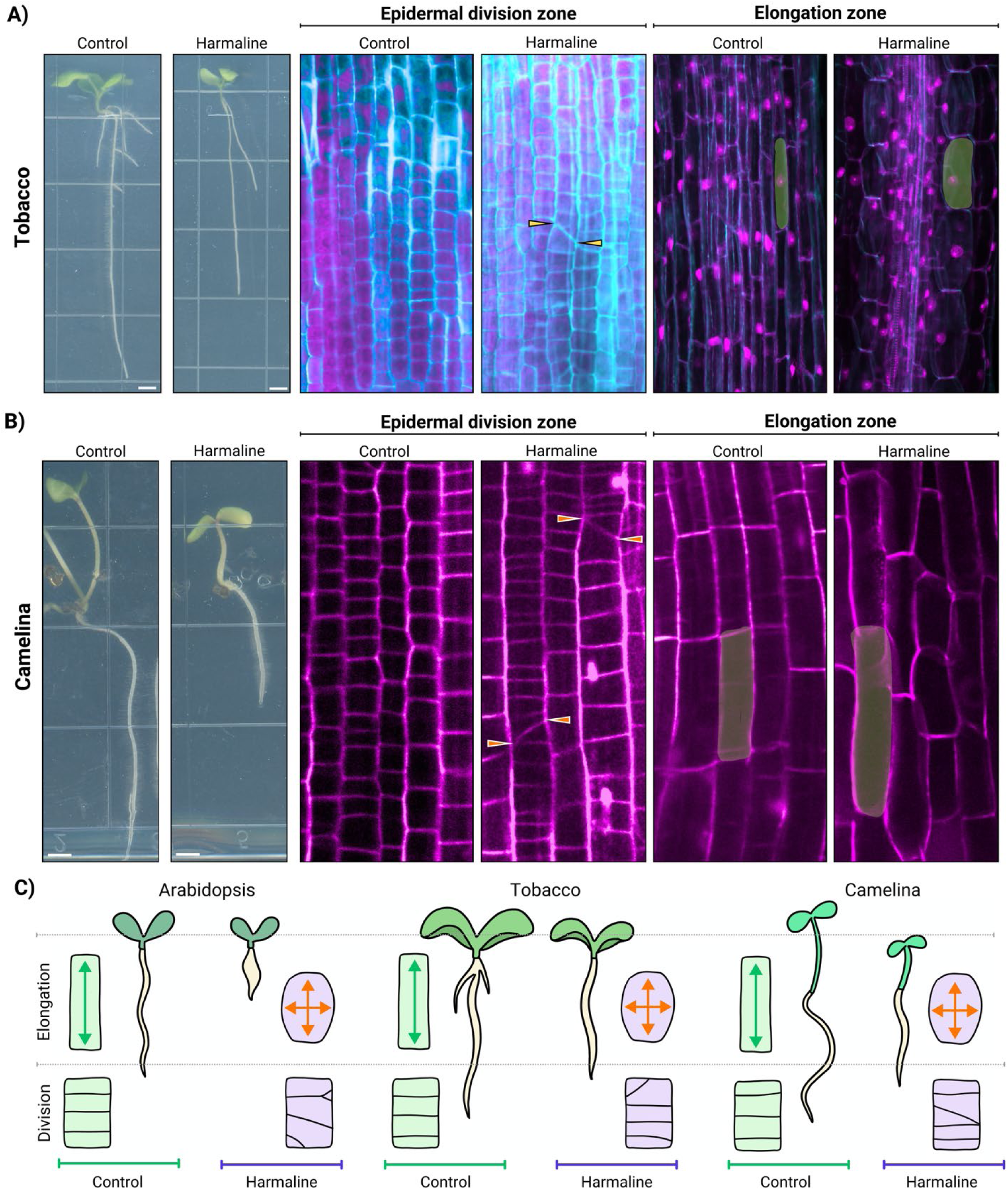
Harmaline induces mild division plane and elongation defects in tobacco and camelina. **A-B)** Representative images of tobacco (A) and camelina (B) seedlings and root tissues grown under control or harmaline treatment conditions. In both species, harmaline induced division plane defects in the epidermal division zone (arrowheads), similar to those observed in Arabidopsis, although the frequency and severity appeared substantially reduced. Likewise, cells in the elongation zone displayed mild radial swelling following harmaline treatment, but this phenotype was less pronounced than that observed in Arabidopsis. C) Model depicting similar phenotypes in three different species.

## Discussion

The harmala alkaloids provide an excellent system for investigating how plants can use a single class of molecules for multiple purposes. Specifically, these alkaloids have been shown to exert anti-feedant properties on insects (Moloudizargari et al., 2013), and previous studies have suggested that they are produced by *P. harmala* for their allelopathic properties (Herraiz et al., 2010). In this study, we found that of the harmala alkaloids, harmaline has the most potent root inhibitory activity and is the only one that causes the distinctive cell division plane defect (Figure 1; Supplemental Figure 2). During the broad search about the potency of harmaline for its allelopathic properties at the tissue and cellular level, we have found that it can inhibit root growth by affecting the cell division plane and cell elongation in a wide range of plant species (Figure 8, 9). To qualify as potent herbicide, selective activity of a molecule would be essential to reduce unwanted plant growth while limiting effects on the crop of interest. The insensitive root growth phenotype of crop plants, including maize, wheat, and tomato suggest a potential use of harmaline, or a molecule with a similar mechanism of action, as an herbicide for agricultural use (Figure 8). However, little was previously known about the cellular mechanism by which harmaline exerts its root inhibition activity. Indeed, root growth inhibition alone provides limited mechanistic insight, as root growth reflects the combined effects of multiple developmental processes including cell division, cell elongation, differentiation, and stress responses. Thus, while root growth inhibition is a useful physiological readout of harmaline activity, the specific cellular phenotype identified here may provide a more informative mechanistic signature of its mode of action.

In this study, we utilized a quantitative live cell imaging approach to understand the cellular and molecular mechanism of harmaline. We found that harmaline-induced cell division defects were highly enriched during late phragmoplast expansion. At first glance, this phenotype resembles a classical phragmoplast guidance defect (Rasmussen et al., 2011; Nan et al., 2023). However, TAN1-YFP remained correctly localized at the cortical division site with normal orientation (Figure 7), indicating that division plane specification and maintenance were not disrupted. Thus, the defect arises despite proper TAN1 positioning, suggesting that harmaline acts downstream of, or independently from, TAN1-mediated division site marking. Together, these findings indicate that harmaline perturbs cell division plane execution rather than initial division plane specification in a TAN1-independent manner. The stage-specific nature of this phenotype also suggests that harmaline does not broadly target microtubules. Cortical microtubules, the preprophase band, spindle structures, and the phragmoplast all depend on microtubule organization, and direct disruption of microtubule dynamics would therefore be expected to produce abundant defects across multiple stages of mitosis. This distinction is particularly noteworthy because many plant-derived alkaloids and alkaloid-derived compounds interfere with microtubule polymerization or dynamics. Classical compounds such as colchicine and vinca alkaloids induce widespread mitotic defects across multiple microtubule arrays rather than highly stage-specific phenotypes (Jordan & Wilson, 2004). In contrast, harmaline predominantly affected late-stage phragmoplast orientation, supporting the possibility that it acts through a mechanism distinct from general microtubule inhibition.

Interestingly, despite the close structural similarity among the harmala alkaloids tested here, harmaline uniquely produced a strong cell division plane defect. Harmine caused only moderate root growth inhibition at concentrations below 50 μg/mL, while neither harmine nor the other tested harmala alkaloids reproduced the distinctive division plane phenotype (Supplementary Figure 2). However, another harmala alkaloid, norharmane, the simplest member of this scaffold class, has also been reported to induce cell division-related defects in Arabidopsis (López-González et al., 2020). These observations emphasize that subtle structural differences within harmala alkaloids may strongly influence biological activity and target specificity. Consistent with this interpretation, transcriptomic analysis provided little evidence that harmaline induced stronger transcriptional responses in cell division-related genes than the structurally similar compound harmine. Despite producing a much more pronounced cell division phenotype, harmaline did not substantially alter expression of known division-associated factors relative to harmine (Figure 8). This disconnect between phenotype severity and transcriptional response suggests that harmaline may not primarily act through large-scale transcriptional reprogramming. Instead, its effects may arise through a more immediate physical or biochemical mechanism, such as perturbation of protein activity, molecular interactions, membrane properties, or subcellular organization.

In this context, harmaline represents a useful chemical-genetic tool to disentangle TAN1-dependent and TAN1-independent mechanisms underlying cell division plane determination. Interestingly, the cell division in plants is maintained by a series of proteins, starting from nuclear position (Ashraf & Facette, 2020; Ashraf et al., 2023; Hazelwood et al., 2025), preprophase band formation (Van Damme et al., 2007; Rasmussen & Bellinger, 2018; Kumari et al., 2021), spindle organization (Müller et al., 2009), phragmoplast orientation (Smertenko et al., 2017), phragmoplast guidance (Müller et al., 2006; Rasmussen et al., 2011), and new cell plate formation (Verma, 2001). A comprehensive reverse genetic approach will provide further understanding of the harmaline-mediated cell division plane determination in plant cells. Furthermore, longer-term time lapse imaging of up to several days will help us to observe a large number of cells going through the full cell division process and capture the harmaline-induced cell division defects in a more precise spatial and temporal manner.

Plant alkaloids represent a vast natural resource for medicines and herbicides, and understanding their biological functions can identify novel mechanisms that plants use to engage with other organisms in their environment. Here, we have shown how *P. harmala* plants may use harmala alkaloids as a mechanism to disrupt normal cell division in other plant species, presumably as a mechanism to out-compete them for resources in the arid ecosystems that they inhabit (Shao et al., 2013). Because harmaline affects cell division machinery, a cellular process that is conserved in multicellular organisms, we anticipate that this mechanism of action may have potential applications in both medicine and agriculture. This study provides an initial step toward defining the function of harmaline at the cellular scale, which will facilitate future studies into the specific molecular target(s) of this alkaloid.

## Acknowledgements

The authors thank Masaaki Umeda, Nara Institute of Science and Technology (Cytrap); Sachihiro Matsunaga, Tokyo University of Science (proPCNA1::PCNA1-sGFP); Abidur Rahman, Iwate University (LTI6b-GFP); Geoffrey Wasteneys, The University of British Columbia (UBQ1pro:GFP-MBD and proTuB2:mScarlet-TuB2); Kenneth Birnbaum, New York University (pro35S::H2B-mRFP1); Carolyn Rasmussen, University of California Riverside (TAN1-YFP); Georgia Drakakaki, University of California Davis (RabA2a-GFP); Arp Schnittger, University of Hamburg (CDKA;1-mVenus/*cdka;1*); Norman Best, United States Department of Agriculture (Maize B73); Michelle Facette, University of Massachusetts Amherst (Maize B73); Liang Song, The University of British Columbia (Camelina, Tomato M82); for sharing seeds.

The authors thank Miki Fujita and EunKyoung Lee of UBC Bioimaging Facility (RRID: SCR_021304) for their kind support on confocal microscope imaging; and Lacey Samuels (The University of British Columbia) and Derrick Horne (Electron Microscopy specialist, UBC Bioimaging Facility) for suggestions and recommendations on performing scanning electron microscopy. RNA sequencing analysis was performed through the Fir-high performance cluster at SFU, enabled by the BC DRI Group and the Digital Research Alliance of Canada. Authors thank Bahman Khahani (University of Massachusetts Amherst) for technical help and suggestion on RNAseq data analysis.

The research at Ashraf Lab is funded by the NSERC Discovery grant (RGPIN-2025-04277), Canada Foundation for Innovation (CFI) John R. Evans Leaders Fund (JELF), BC Knowledge Development Fund (BCKDF), and start-up grant provided by the University of British Columbia. Olivia S. Hazelwood received NSERC CGRS-D fellowship, the 4-year doctoral fellowship by the University of British Columbia, Linda Matsuuchi award, Hugo E Meilicke Memorial Fellowship, and Frances Chave Memorial Graduate Scholarship. Joh Demura-Devore received the UBC’s work-learn international undergraduate research fellowship.

## Author contributions

SOD performed the initial experiments in identifying the distinct roles of harmala alkaloids and cell division plane defects caused by harmaline. KAD performed the time course and live cell imaging experiment and quantification of confocal microscopy. OSH performed the scanning electron microscopy experiment. SOD and OSH set up the experiment and isolated RNA samples for sequencing. KAD and JD performed the RNAseq data analysis. RSN, FM, and MAA adopted the initial project idea. KAD and MAA designed the quantitative image analysis parameters, prepared the figures, and performed the statistical analysis. KAD, FM, RSN, and MAA wrote the final version of the manuscript and all the authors read and approved it. MAA acquired the funding and supervised KAD, SOD, OSH, and JD.

## Data availability

The data that support the findings of this study are openly available at E-MTAB-17129 in the European Nucleotide Archive.

---

**Supplemental Figure 1:**
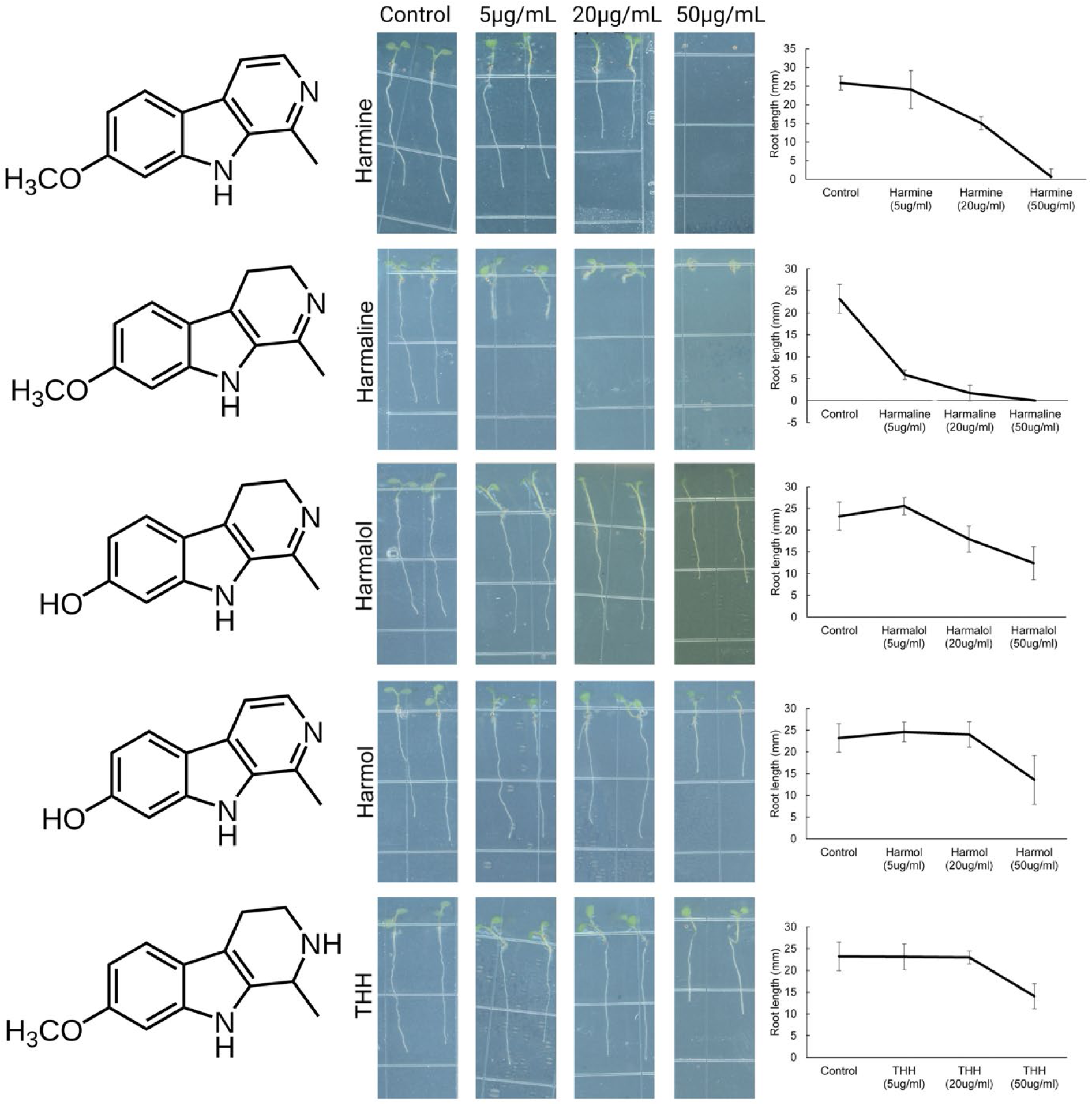
Dose response curve for Harmine, Harmaline, Harmalol, Harmol, and Tetrahydroharmine. The chemical structure (left panel), root growth phenotype (middle panel) of 5 days old seedlings (Control, 5, 20, and 50 mg/ml concentration), and quantification (right panel) of Harmine, Harmaline, Harmalol, Harmol, and Tetrahydroharmine (top to bottom).

**Supplemental Figure 2.**
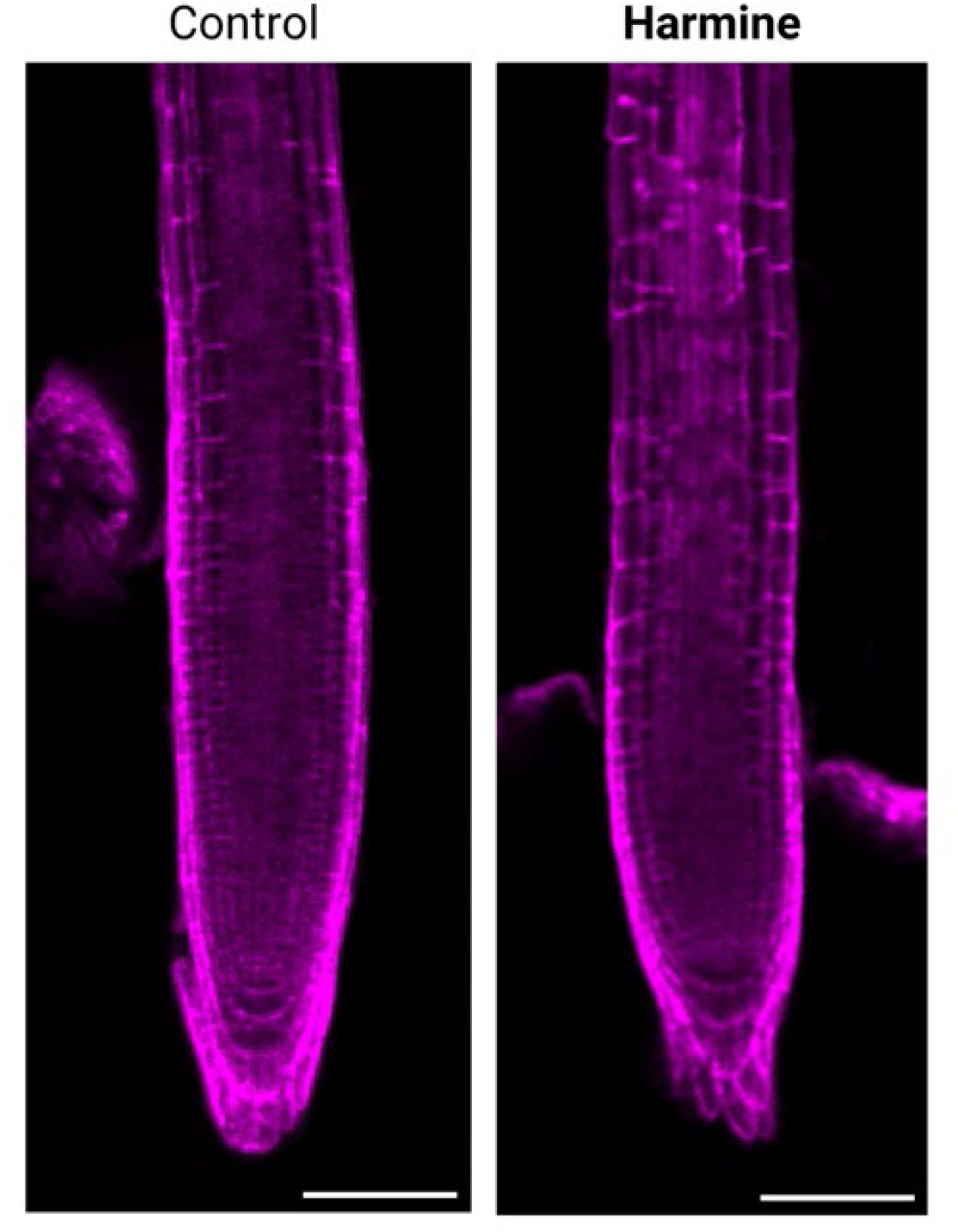
Representative confocal images of Arabidopsis primary roots treated with harmine. Although harmine was the second most potent harmala alkaloid tested for inhibition of root growth, treated roots did not exhibit the pronounced cell swelling or obvious cell division plane defects observed following harmaline treatment. Scale bars = 100 μm.

**Supplemental Figure 3:**
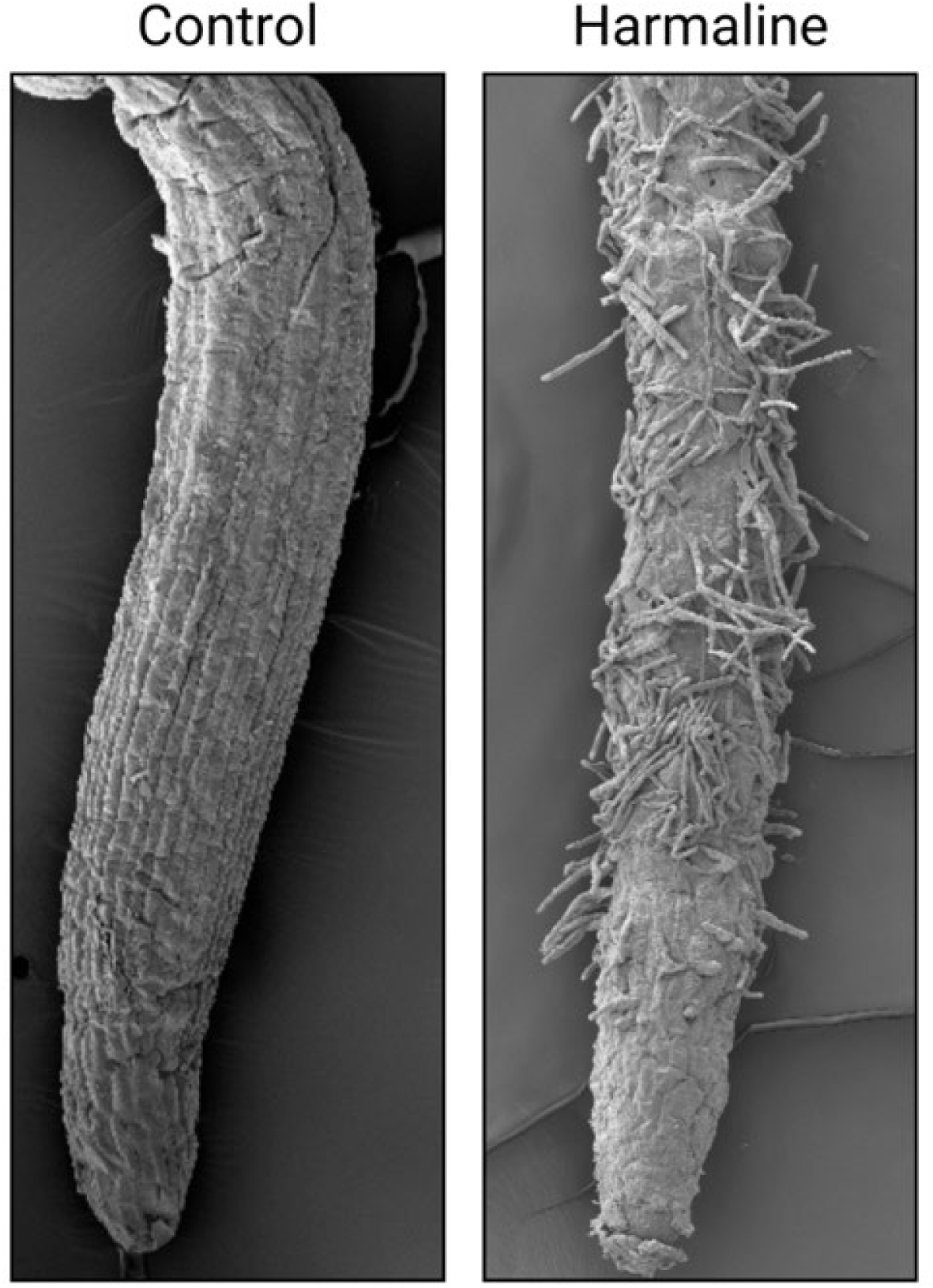
Scanning electron microscopy (SEM) of Arabidopsis roots under control and harmaline treatment. Representative SEM images show primary root morphology in control (left) and harmaline-treated (right) seedlings. In control roots, root hairs emerge in the expected differentiation zone located proximally from the root tip. In contrast, harmaline-treated roots exhibit a clear shift in root hair positioning, with abundant root hair formation occurring closer to the root tip.

**Supplemental figure 4.**
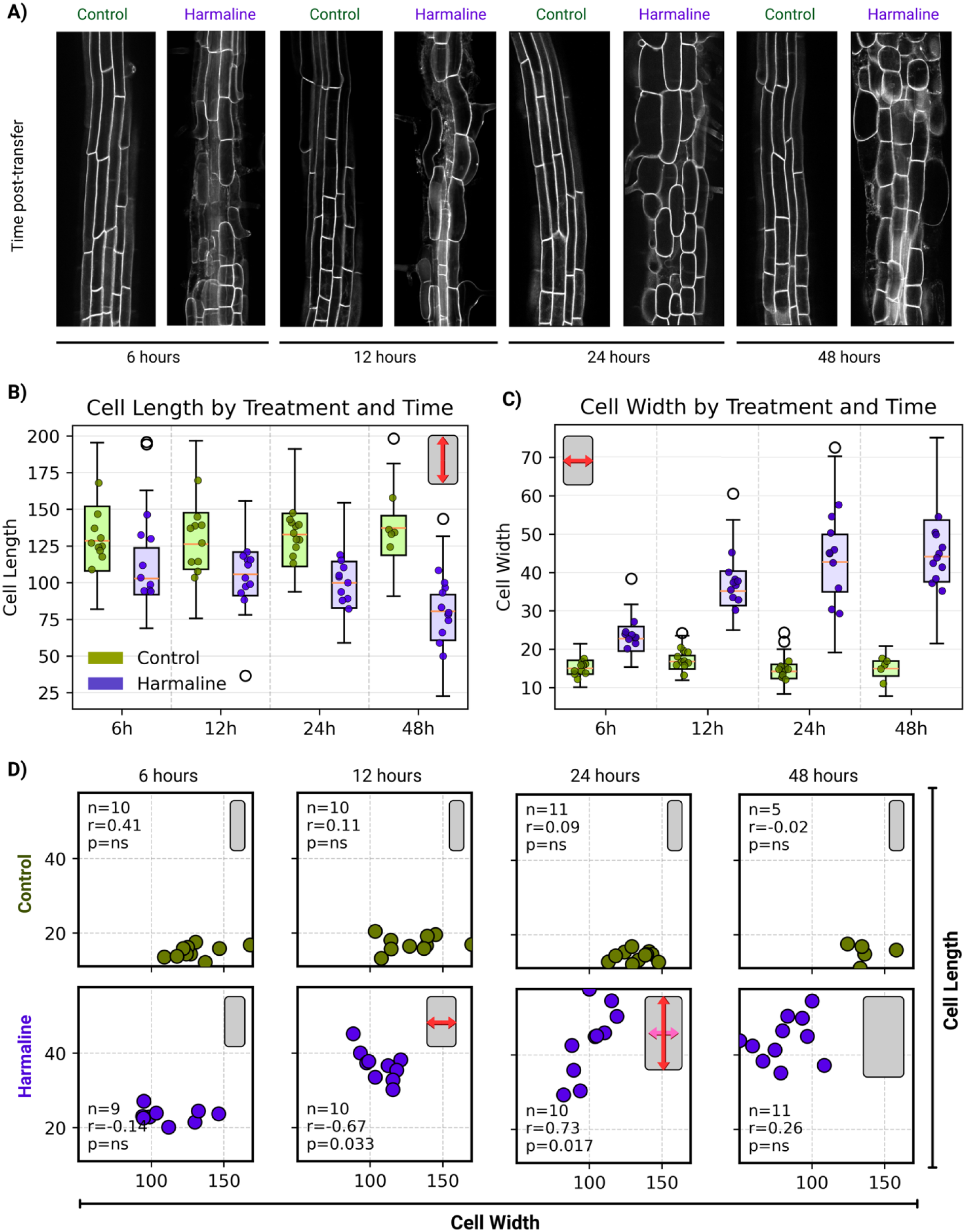
Harmaline induces rapid radial swelling of elongating root cells. **A)** Representative confocal images of elongation zone cells in seedlings transferred to control or harmaline-containing media and imaged at 6, 12, 24, and 48 hours post-transfer. Control cells maintain elongated morphology, whereas harmaline-treated cells show progressive radial expansion and loss of anisotropic growth over time. **B)** Quantification of cell length across timepoints. Harmaline-treated cells are significantly shorter than controls at all timepoints. Mean cell lengths (± SD) and sample sizes were: 6 h, Control 128.5 ± 28.5 µm (n = 48) vs Harmaline 102.9 ± 26.8 µm (n = 45), p = 7.67 × 10⁻⁴; 12 h, Control 127.7 ± 21.8 µm (n = 46) vs Harmaline 96.0 ± 23.5 µm (n = 73), p = 2.40 × 10⁻⁶; 24 h, Control 138.3 ± 22.7 µm (n = 73) vs Harmaline 96.5 ± 22.5 µm (n = 84), p = 3.39 × 10⁻⁶; 48 h, Control 137.2 ± 26.5 µm (n = 29) vs Harmaline 80.5 ± 23.8 µm (n = 67), p = 3.67 × 10⁻¹³. **C)** Quantification of cell width across timepoints. Harmaline treatment causes a significant increase in cell width at all timepoints. Mean cell widths (± SD) and sample sizes were: 6 h, Control 15.0 ± 2.5 µm (n = 48) vs Harmaline 22.8 ± 4.5 µm (n = 45), p = 1.81 × 10⁻¹⁵; 12 h, Control 17.4 ± 2.9 µm (n = 46) vs Harmaline 36.7 ± 8.6 µm (n = 73), p = 1.06 × 10⁻²³; 24 h, Control 14.7 ± 2.8 µm (n = 73) vs Harmaline 42.7 ± 11.9 µm (n = 84), p = 7.33 × 10⁻⁸; 48 h, Control 14.5 ± 3.1 µm (n = 29) vs Harmaline 44.1 ± 11.5 µm (n = 67), p = 3.64 × 10⁻³⁴. In boxplots, boxes represent the interquartile range, the central line indicates the median, whiskers extend to 1.5× the interquartile range, and dots indicate the mean of individual replicates. Statistical comparisons were performed using Welch’s t-tests on pooled cell measurements**. D)** Schematic representations of cell geometry corresponding to the observed correlations between cell width and cell length across time points (6, 12, 24, and 48 hours) in control and harmaline-treated samples. In control conditions, cell shape remains consistent over time, with no clear relationship between width and length. In contrast, harmaline treatment induces dynamic changes in cell geometry. At 12 hours, cells display a negative relationship between width and length, represented by wider but shorter cells. At 24 hours, this shifts to a positive relationship, with cells increasing in both width and length. By 48 hours, this relationship weakens, and cell proportions return to a more variable, non-correlated state. These schematics summarize the temporal disruption of coordinated cell growth under harmaline treatment.

**Supplementary figure 5.**
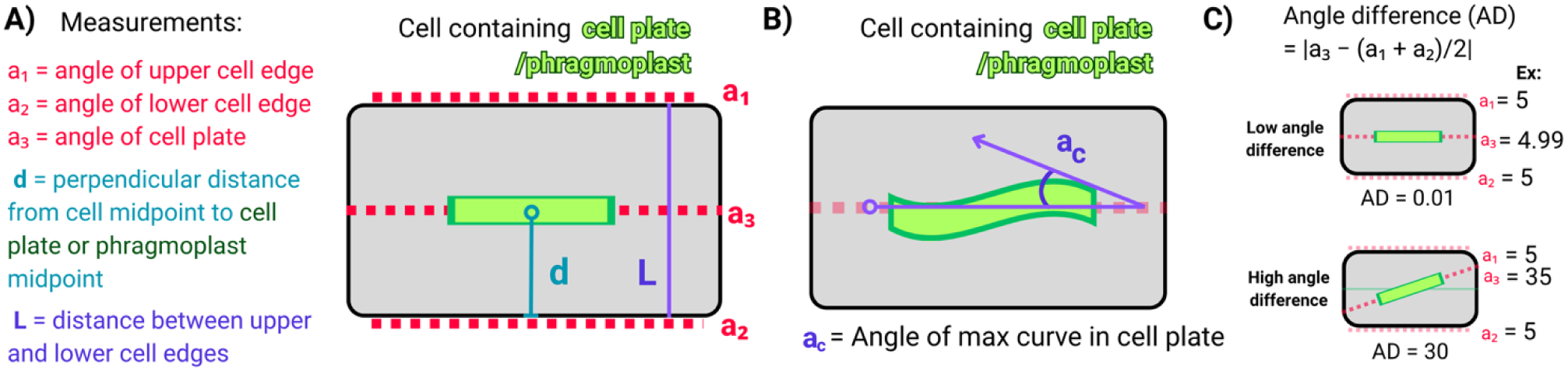
Quantification of phragmoplast and cell plate geometry. **A)** Schematic of measurements used to quantify cell plate and phragmoplast positioning and orientation within dividing plant cells. The angles of the upper (a₁) and lower (a₂) cell edges were defined relative to the longitudinal cell axis, and the angle of the cell plate (a₃) was measured at the plane of division. The perpendicular distance (d) from the cell midpoint to the midpoint of the cell plate or phragmoplast was used to assess positional deviation. The distance between the upper and lower cell edges (L) defines the cell width along the measurement axis. Measurements were performed on either the cell plate or phragmoplast midzone, depending on visibility. **B)** Measurement of curvature within the cell plate. The angle of maximum curvature (a_c) was measured using the angle tool in ImageJ/Fiji. A reference line was drawn along the midplane that the phragmoplast is expected to follow, and the curvature angle was defined at the point of maximum deviation (top of the curve) relative to this reference. **C)** Quantification of alignment between the cell plate and the cell axis using angle difference (AD), calculated as AD = |a₃ − (a₁ + a₂)/2|. Low AD values indicate proper alignment of the cell plate with the division axis, whereas high AD values indicate misalignment.

**Supplementary Figure 6.**
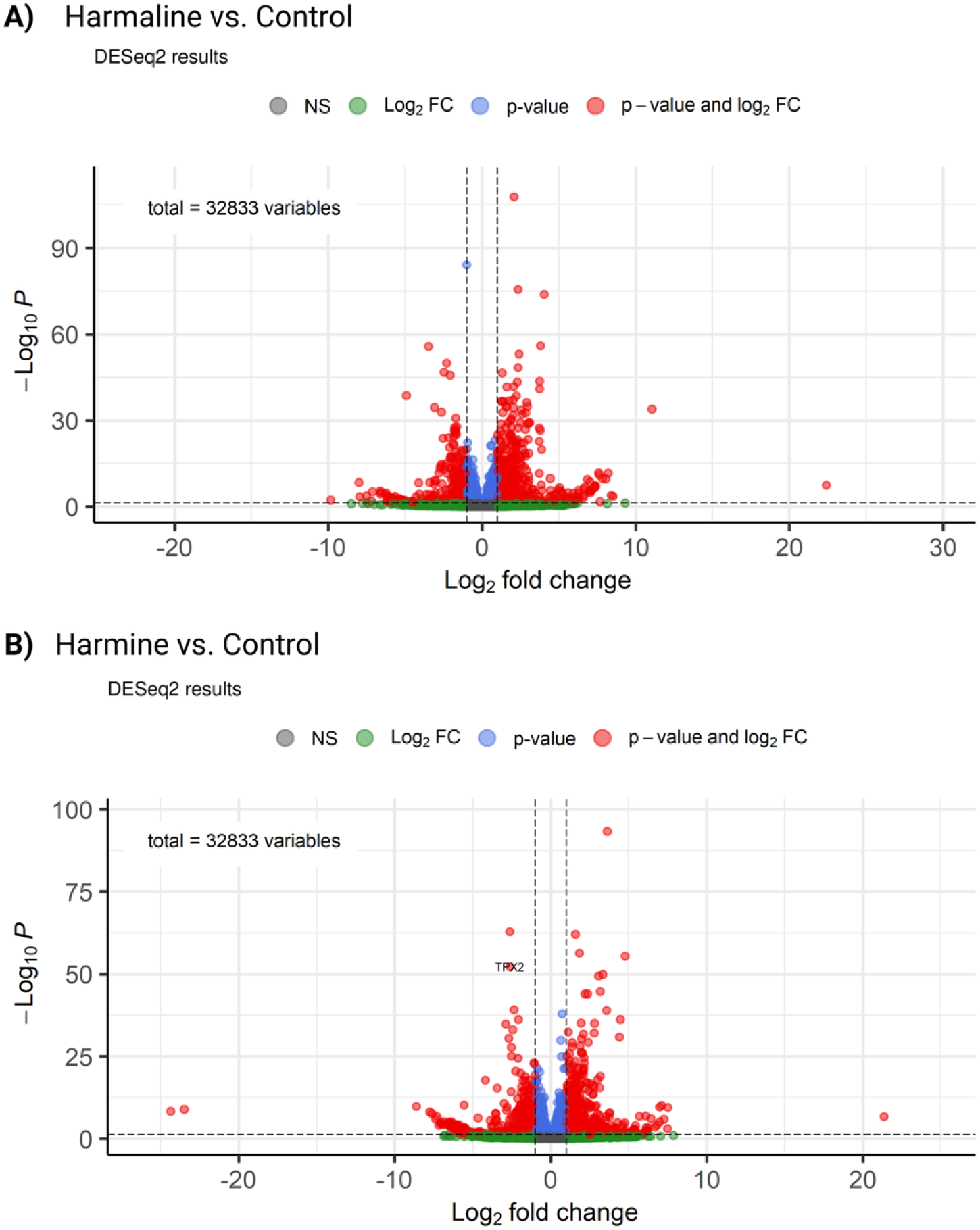
Transcriptomic comparison of harmaline- and harmine-treated Arabidopsis seedlings. **A-B)** Volcano plots showing differentially expressed genes identified by DESeq2 analysis following harmaline **(A)** or harmine **(B)** treatment relative to control samples.

**Supplementary Figure 7.**
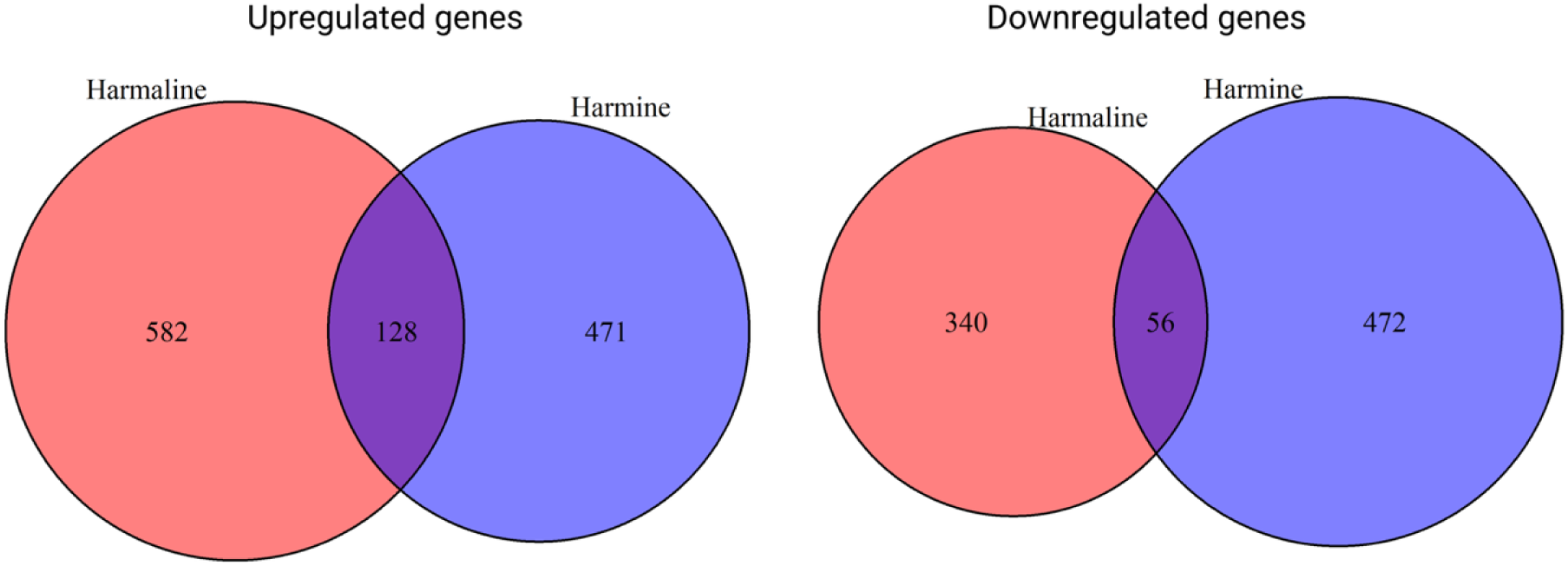
Venn diagrams showing overlap and unique sets of upregulated and downregulated genes between harmaline and harmine treatments.

**Supplementary Figure 8.**
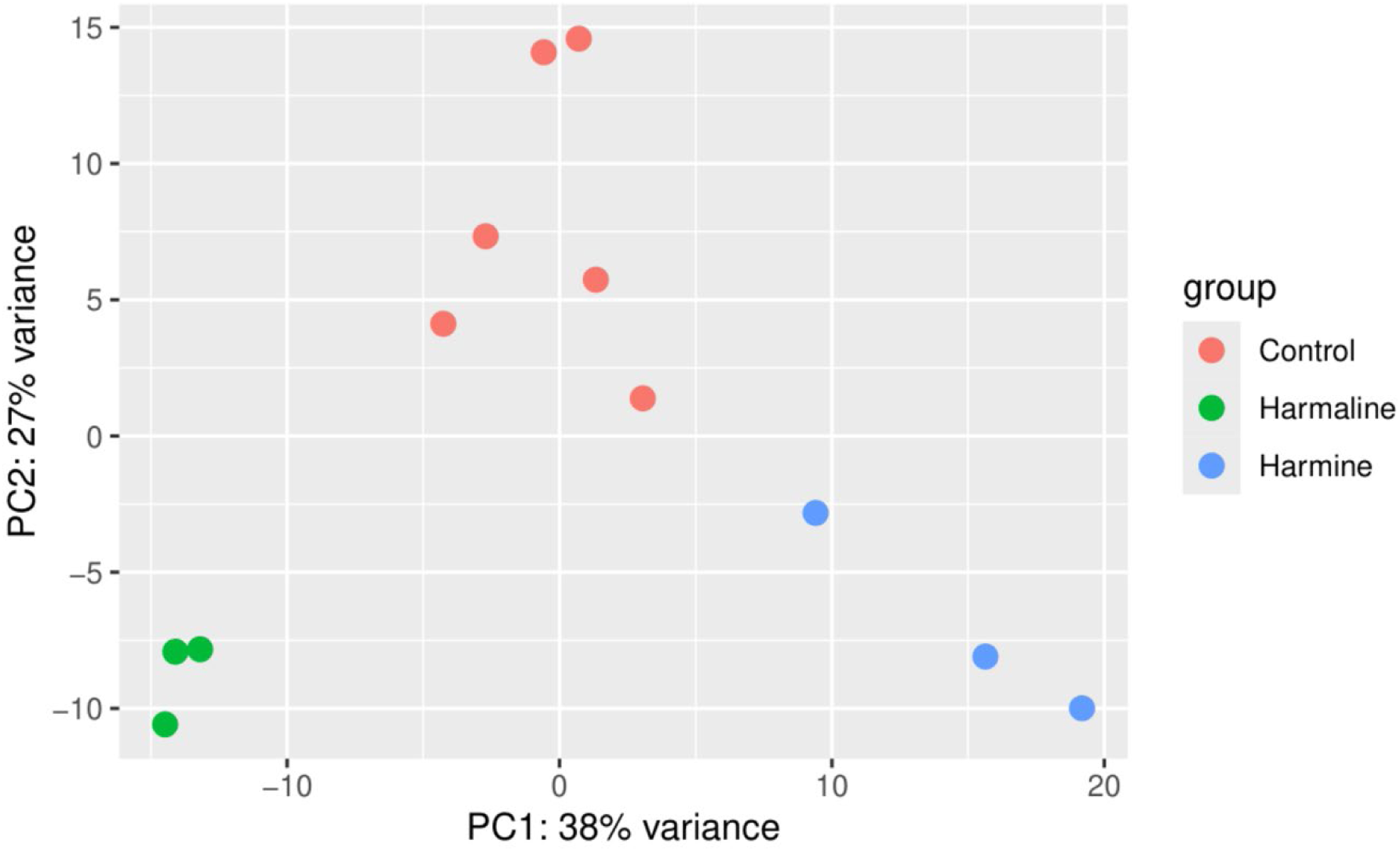
Principal component analysis (PCA) of transcriptomic profiles demonstrating separation among treatment groups.

**Supplementary Figure 9.**
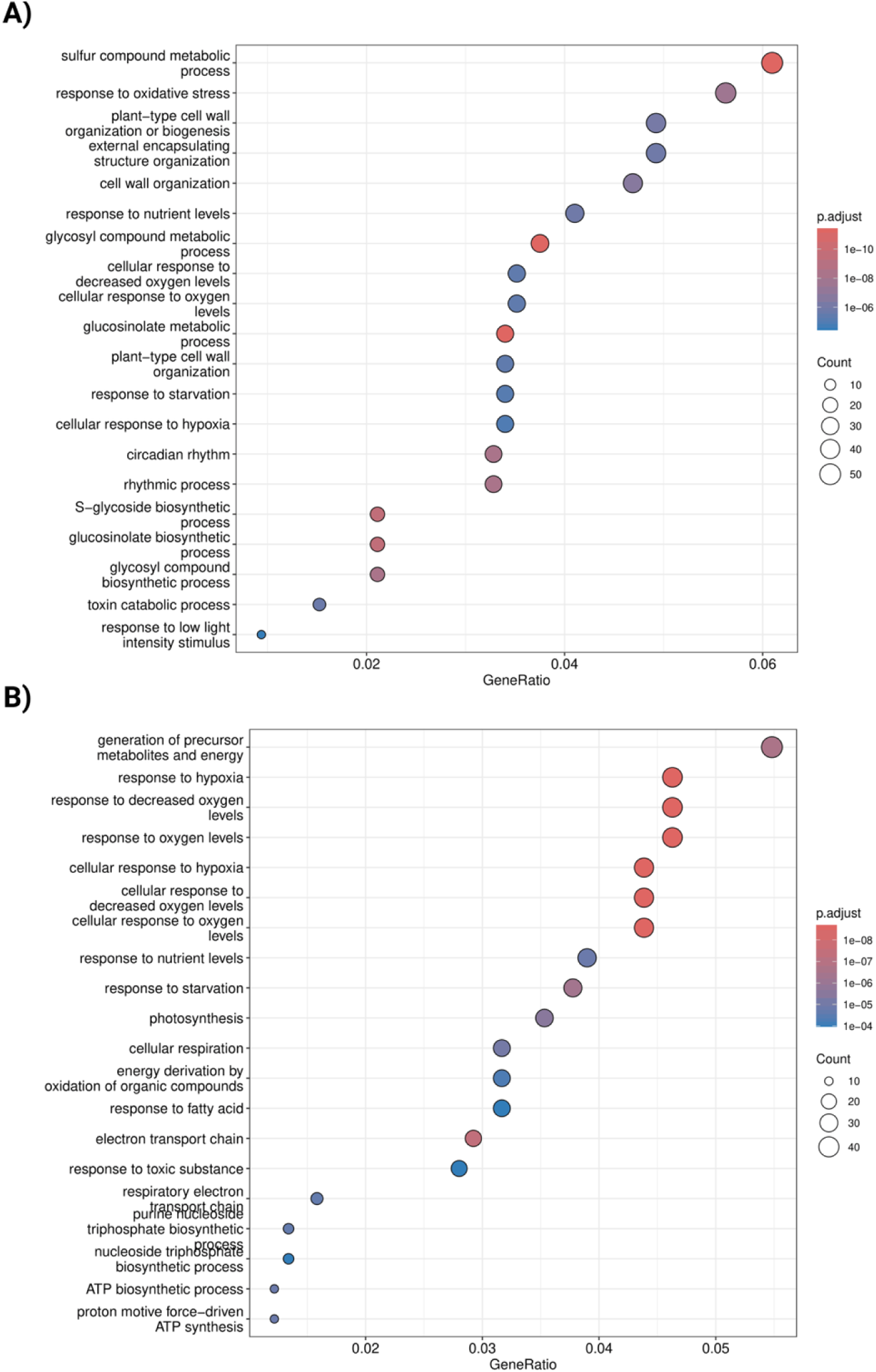
Gene ontology enrichment analysis of differentially expressed genes associated with harmaline (A) and harmine (B) treatments, highlighting enriched biological processes.

## Supplemental videos

**Supplemental Video 1. Time-lapse visualization of Arabidopsis root growth under control and harmaline treatment conditions.** Representative GIF showing seedlings grown on 1/2 MS medium supplemented with 1% sucrose in the presence or absence of 3 μg/mL harmaline. Seedlings were grown under continuous light conditions for 6 days post transfer 2 DAG to visualize overall root growth dynamics and treatment-associated effects on development.

**Supplemental Video 2. Rotating visualization of Arabidopsis root structure with LTi6b-GFP plasma membrane marker.** Representative rotating three-dimensional reconstruction of control and harmaline-treated Arabidopsis roots showing overall tissue organization and root morphology. Seedlings were grown on 1% 1/2 MS media for 2 days and then transferred onto the same media plates with or without 3ug/mL harmaline. Seedlings were imaged 48 hours post-transfer.

**Supplemental Video 3. Live visualization of cell division dynamics in control and harmaline-treated roots.** Representative time-lapse imaging of Arabidopsis root cells expressing GFP-MBD (microtubule binding domain) and H2B-RFP (histone H2B) under control conditions or following treatment with 3 μg/mL harmaline. Seedlings were grown on 1/2 MS medium supplemented with 1% sucrose for 2 days and transferred to the same medium with or without harmaline. Imaging was performed 48 hours post-transfer to visualize cell division dynamics over time.

**Supplemental Video 4. Rotational three-dimensional visualization of Arabidopsis root architecture under control and harmaline treatment conditions.**

Representative rotating three-dimensional reconstruction of Arabidopsis root cells expressing GFP-MBD x H2B-RFP under control conditions or following treatment with 3 μg/mL harmaline. Seedlings were grown on 1/2 MS medium supplemented with 1% sucrose for 2 days and transferred to the same medium with or without harmaline. Imaging was performed 48 hours post-transfer to visualize overall root organization and morphology.

**Supplemental Video 5. Comparison of Z-projection and three-dimensional reconstruction of dividing Arabidopsis root cells.** Representative visualization of Arabidopsis root cells expressing GFP-MBD x H2B-RFP under control and 3 μg/mL harmaline treatment conditions. The movie compares conventional Z-projection and three-dimensional reconstruction approaches to better show how the harmaline-treated phragmoplast is being affected.

**Supplemental Video 6. Visualization of cell plate morphology in harmaline-treated Arabidopsis roots.** Representative live-cell imaging of Arabidopsis roots expressing RabF2a-GFP with propidium iodide staining following treatment with 3 μg/mL harmaline. The movie highlights cell plate organization and morphology, including representative examples of irregular and wavy cell plate structures.

**Supplemental Video 7. Z-stack visualization of Arabidopsis root organization under control and harmaline treatment conditions.** Representative Z-stack imaging of Arabidopsis roots expressing RabF2a-GFP with propidium iodide staining under control conditions or following treatment with 3 μg/mL harmaline. Sequential optical sections through the root are shown to visualize cellular organization and morphology throughout the tissue.

**Supplemental Video 8. Time-lapse visualization of Maize root growth under control and harmaline treatment conditions.** Germinated maize seedlings were grown with and without harmaline for 3 days to observe the root growth.

**Supplemental Video 9. Time-lapse visualization of Wheat root growth under control and harmaline treatment conditions.** Germinated wheat seedlings were grown with and without harmaline for 3 days to observe the root growth.

**Supplemental Video 10. Time-lapse visualization of Tobacco root growth under control and harmaline treatment conditions.** Germinated tobacco seedlings were grown with and without harmaline for 7 days to observe the root growth.

## Supplemental Tables

**Supplemental table 1.** Volcano Plot Data Harmaline vs Control. **Supplemental table 2.** Volcano Plot Data Harmine vs Control.

**Supplemental table 3.** Venn Diagram Data Harmaline and Harmine. **Supplemental table 4.** PCA plot of Harmaline and Harmine.

**Supplemental table 5.** GO dotplot of Harmaline vs Control.

**Supplemental table 6.** GO dotplot of Harmine vs Control.

