## Supplementary figures and images for "Harmala alkaloids regulate cell division planes in plants"

### Supplemental figure 1.tiff

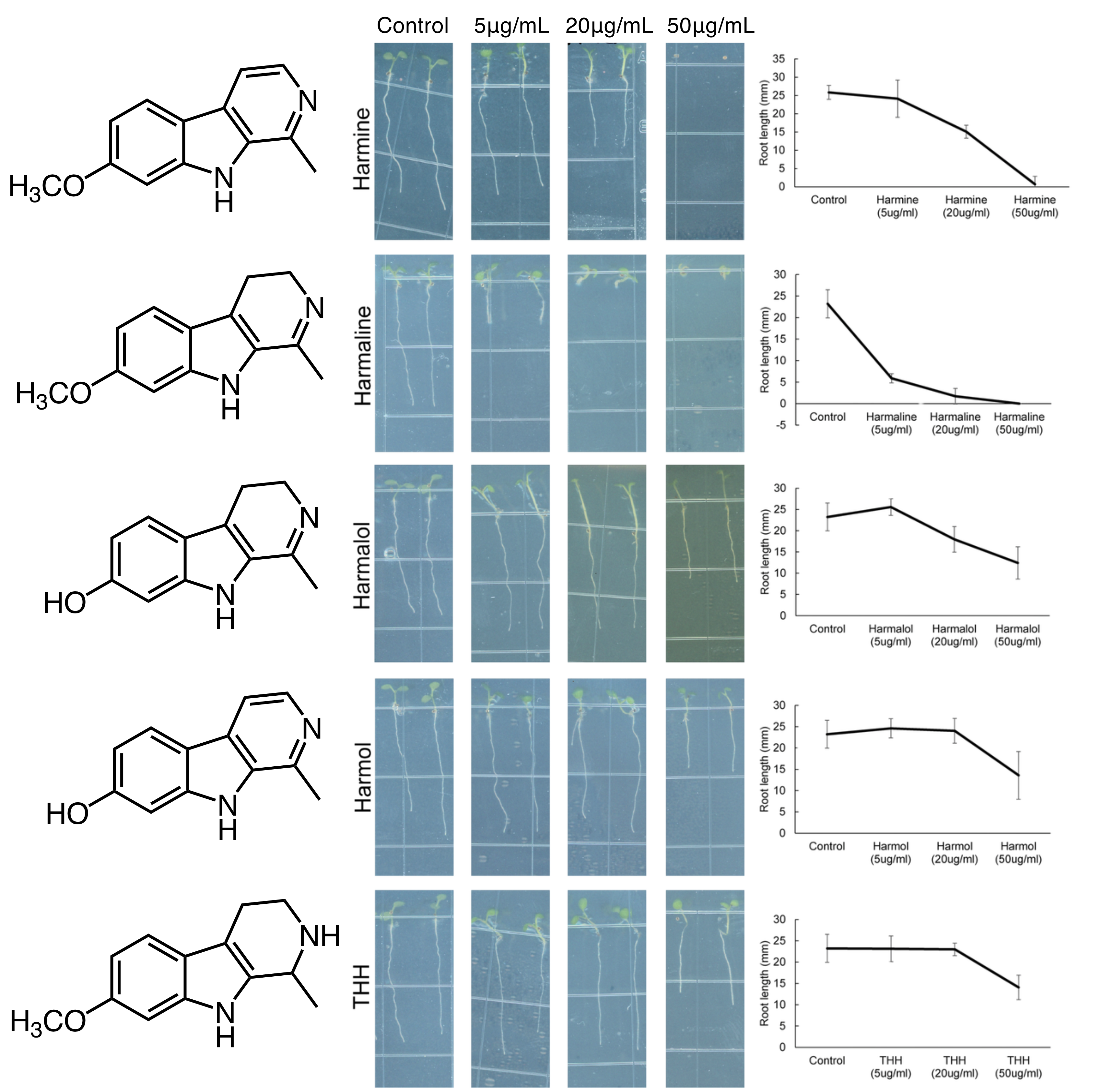

### Supplemental figure 2.tiff

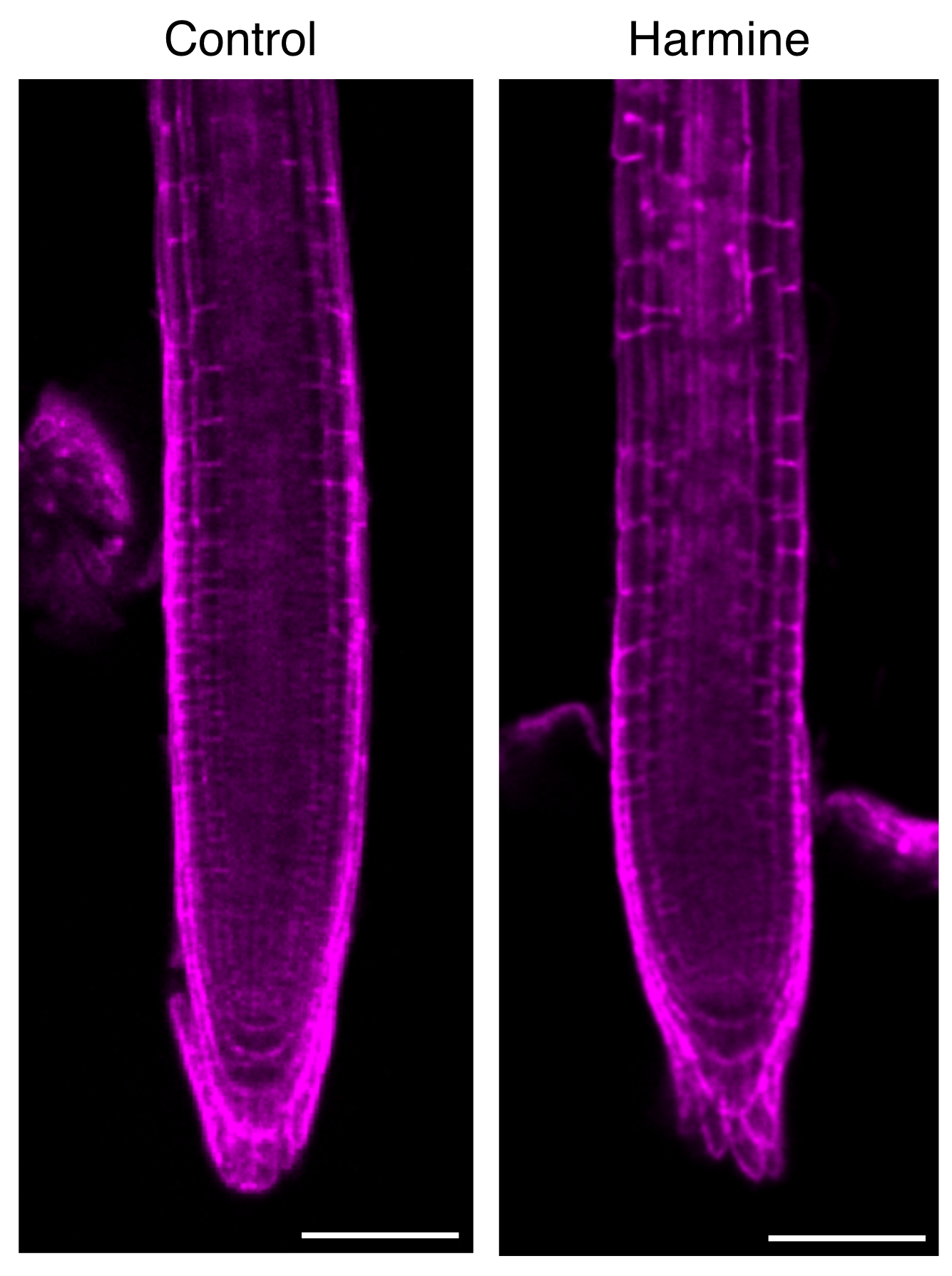

### Supplemental figure 3.tiff

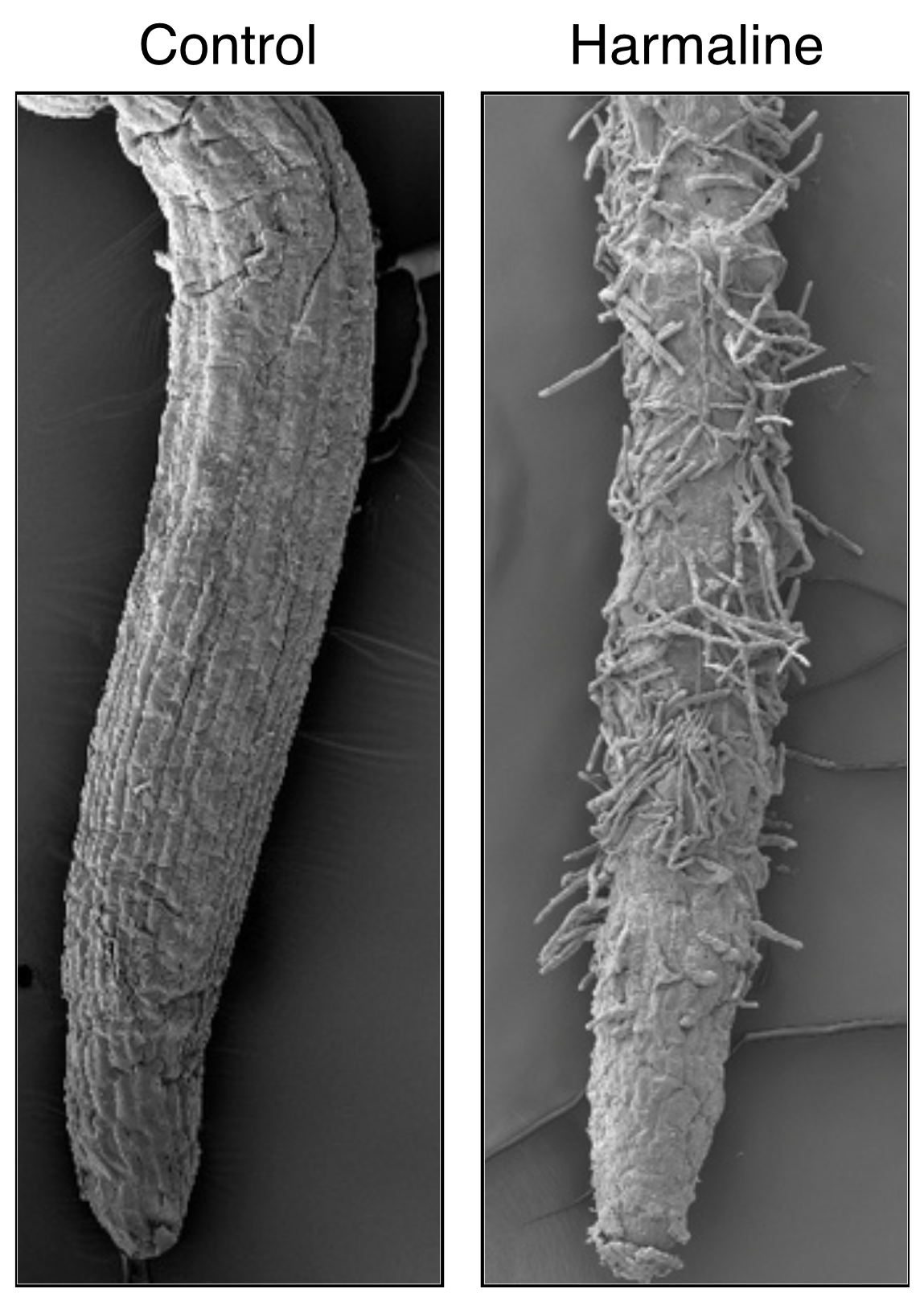

### Supplemental figure 4.tiff

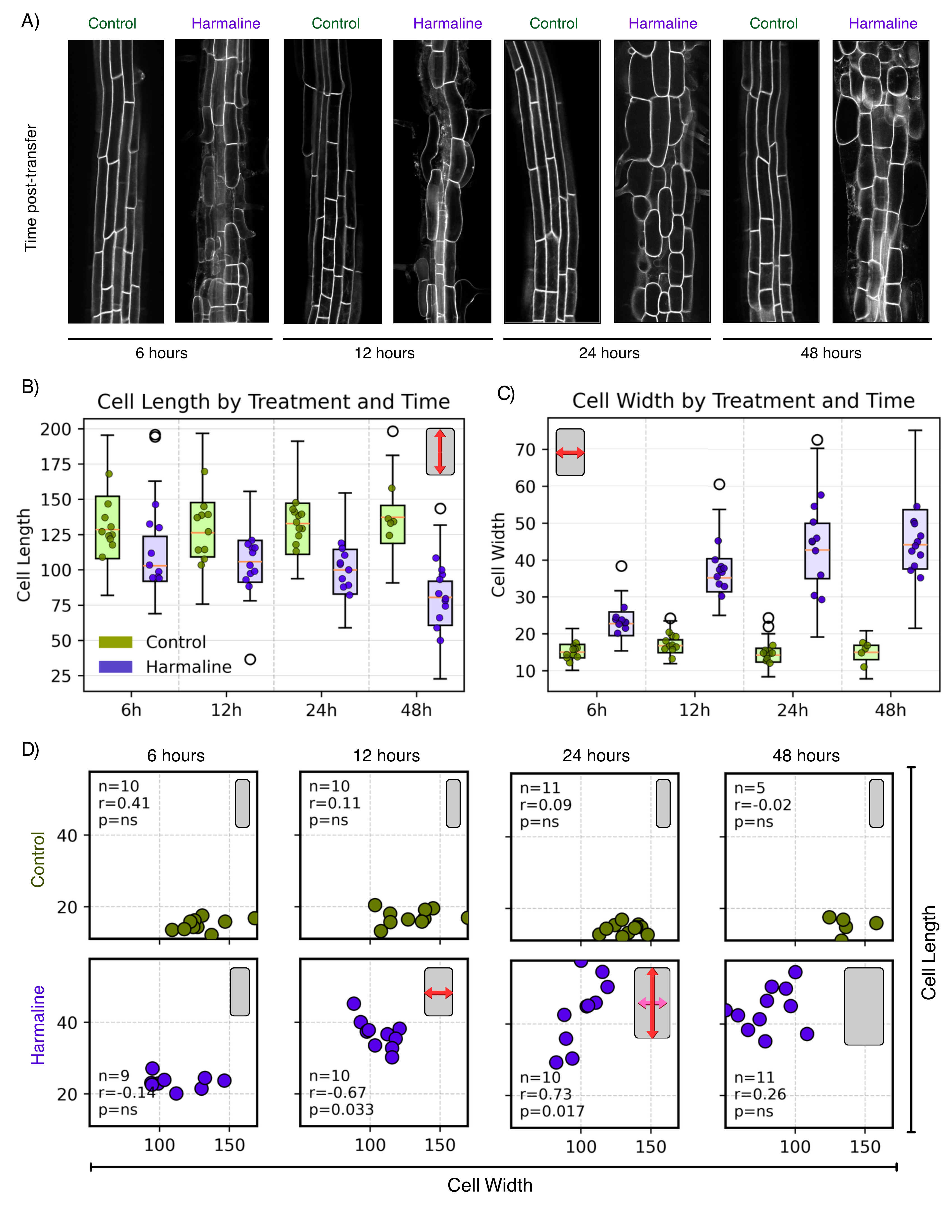

### Supplemental figure 5.tiff

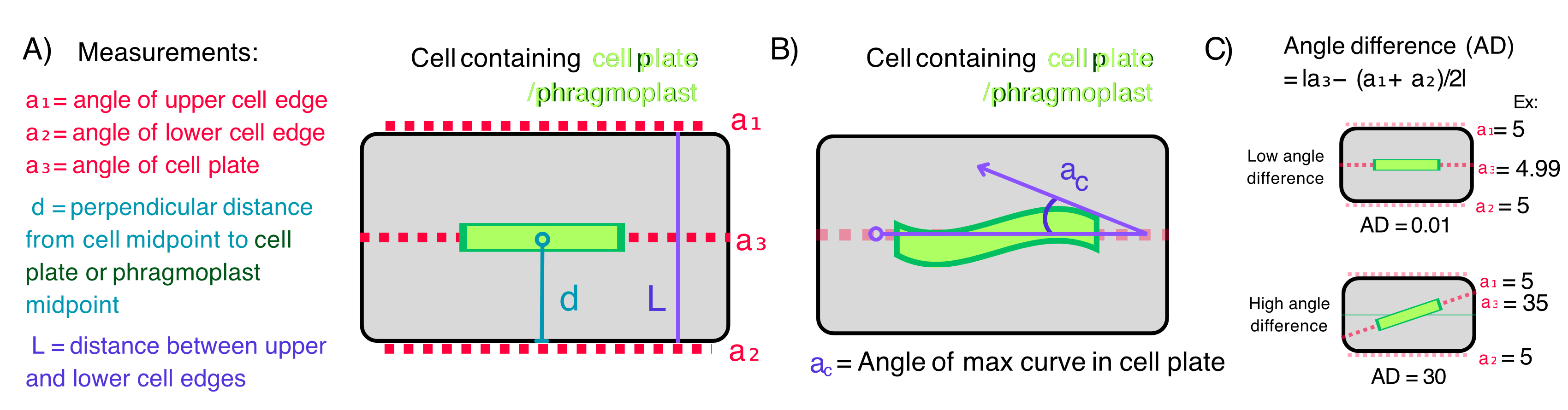

### Supplemental figure 6.tiff

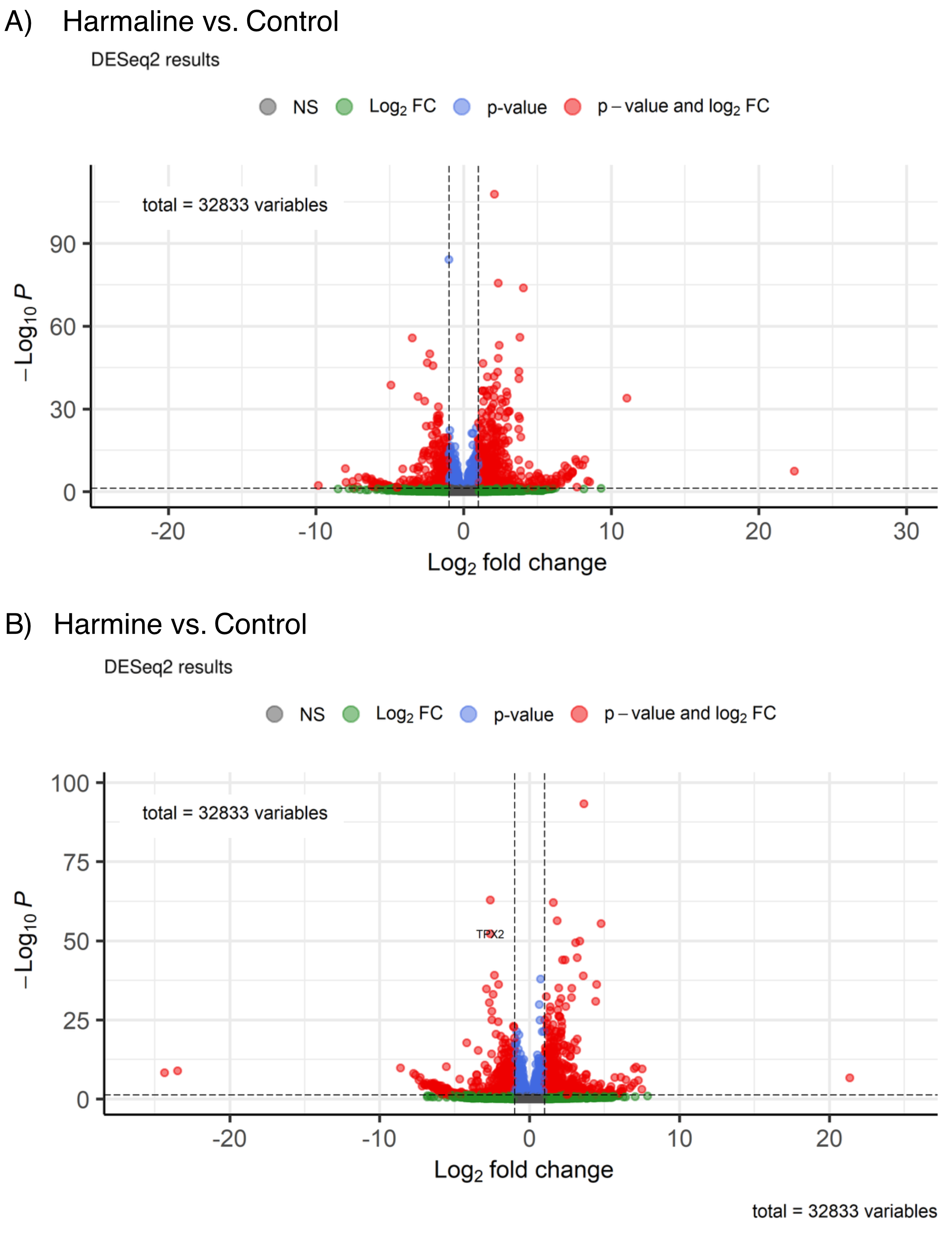

### Supplemental figure 7.tiff

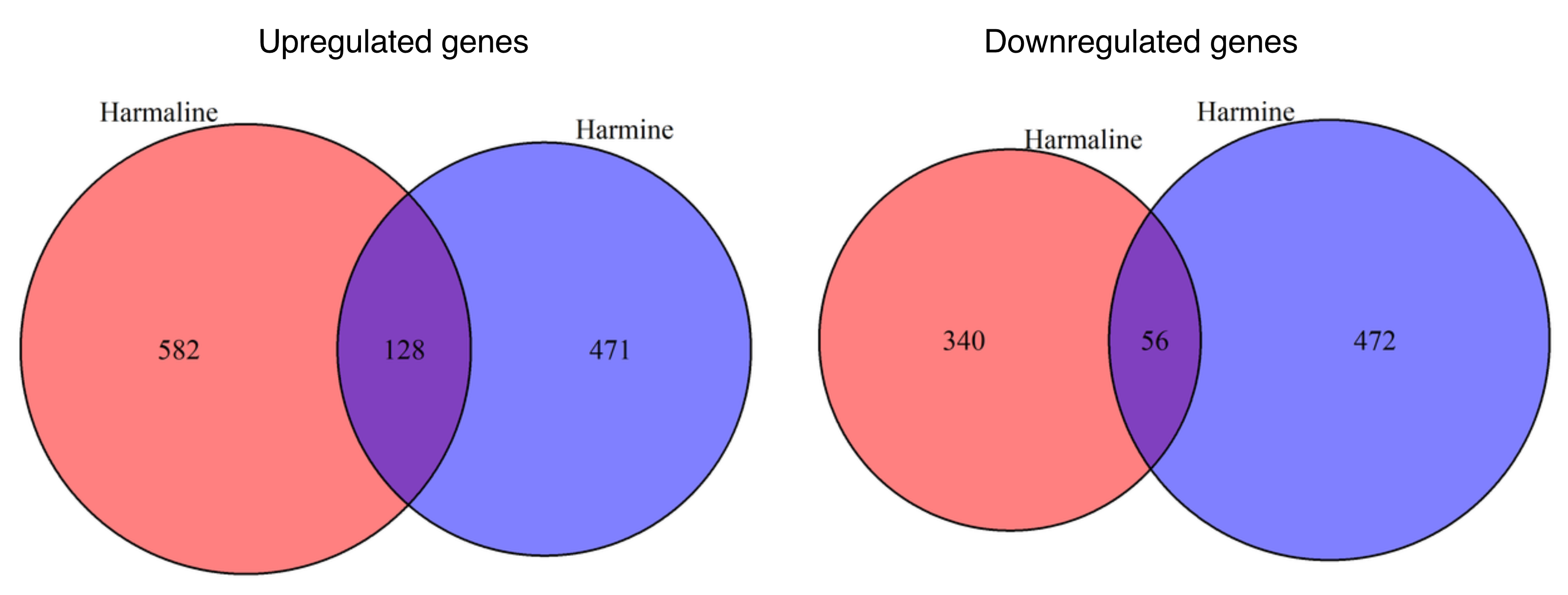

### Supplemental figure 8.tiff

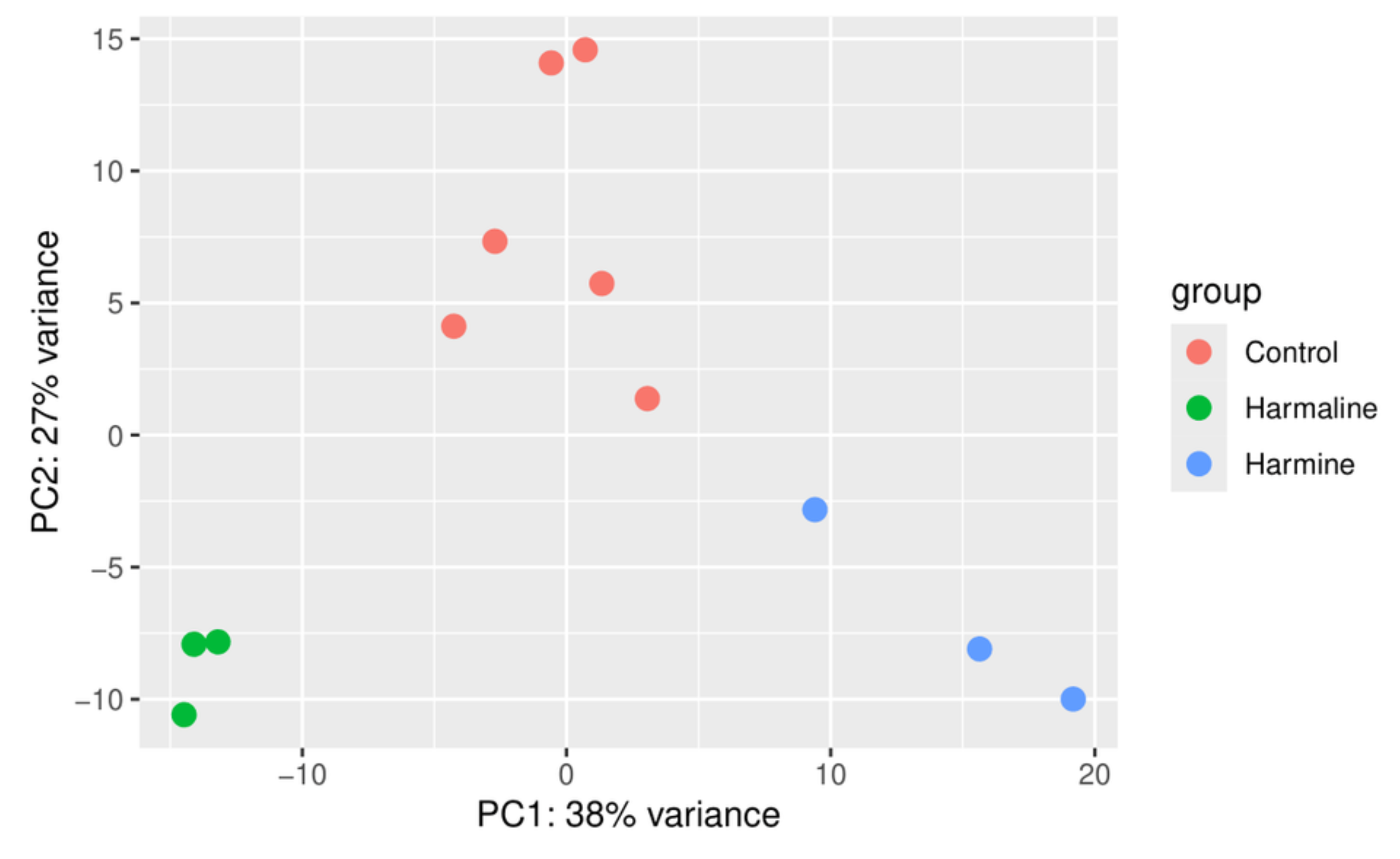

### Supplemental figure 9.tiff

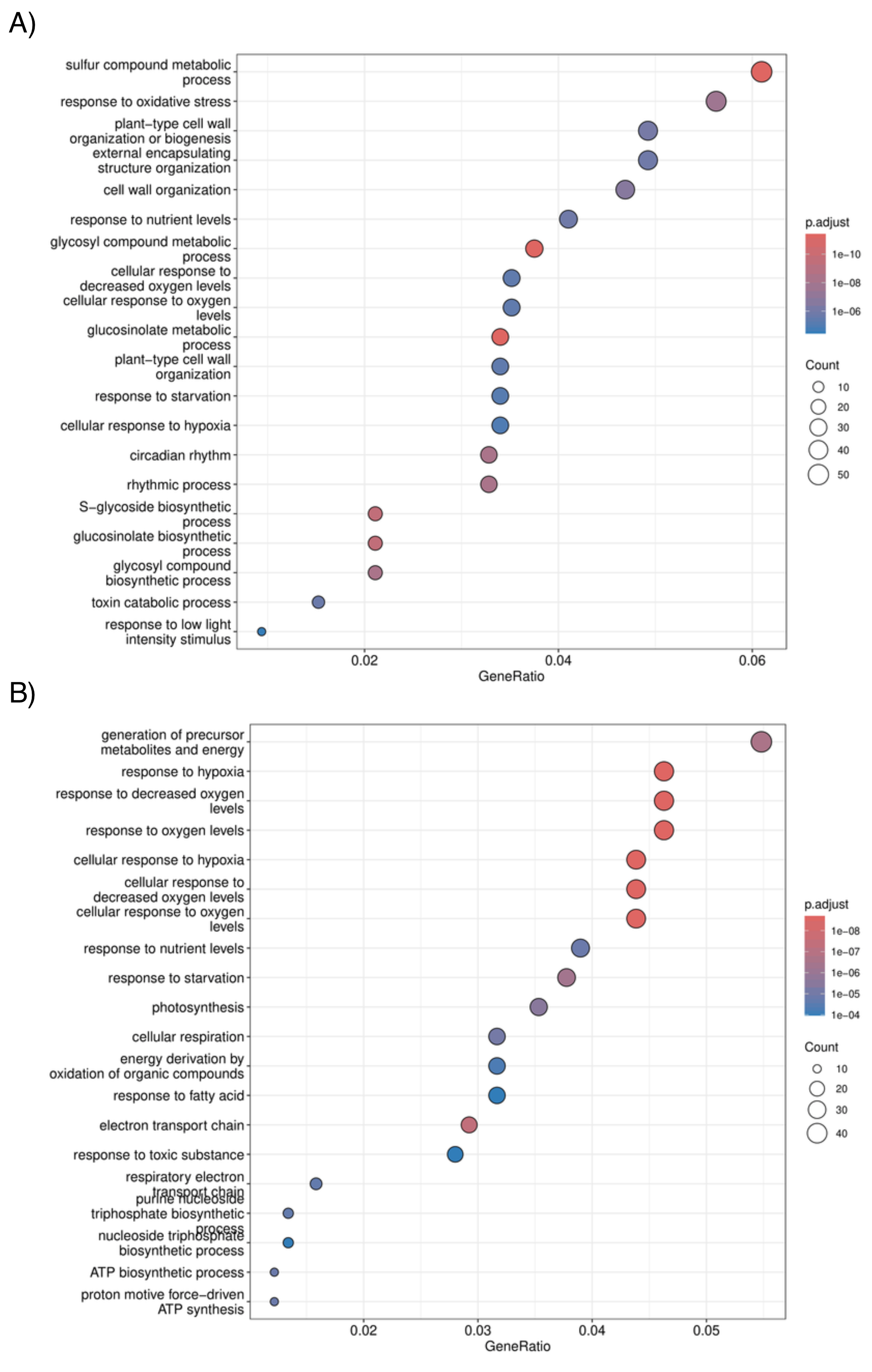
